# Drug treatment alters performance in a neural microphysiological system of information processing

**DOI:** 10.1101/2025.01.27.635163

**Authors:** Bradley Watmuff, Forough Habibollahi, Candice Desouza, Moein Khajehnejad, Alon Loeffler, Koby Baranes, Noah Poulin, Mark Kotter, Brett J. Kagan

**Affiliations:** Cortical Labs Pte. Ltd., Melbourne, VIC, Australia; Turner Institute for Brain and Mental Health, Monash University, Clayton, 3800, VIC, Australia; Department of Clinical Neurosciences, University of Cambridge, Cambridge CB2 0QQ, United Kingdom; Wellcome-MRC Cambridge Stem Cell Institute, University of Cambridge, Cambridge CB2 0AW, United Kingdom; Bit Bio LTD, Cambridge, United Kingdom; Department of Biochemistry and Pharmacology, University of Melbourne, Parkville, VIC, Australia

**Keywords:** iPSC, MEA, electrophysiology, disease model, neuroscience, drug discovery

## Abstract

Assessment of pharmacological intervention on *in vitro* neural systems often emphasizes molecular and structural changes. However, neural systems fundamentally process and act on information. For preclinical assays to predict drug efficacy, they must model these physiological functions. *DishBrain*, an *in vitro* synthetic biological intelligence (SBI) assay embodying a neural system in a simulated game-world, enables the quantification of this information-processing capacity, however the question remains whether such a system permits classical pharmacological interrogation and dose-response profiling. Hyperactive glutamatergic dysregulation is linked to neurological disorders including epilepsy, and inducible overexpression of neurogenin 2 (NGN2) in human induced pluripotent stem cells (hiPSCs) generates glutamatergic cultures with dysregulated hyperactivity. We therefore tested three anti-seizure medications (ASMs), phenytoin, perampanel, and carbamazepine, on NGN2 neurons from day 21 of differentiation in this system. The key finding was that, while all compounds altered spontaneous firing, carbamazepine 200 *µ*M significantly improved gameplay metrics. This marks the first demonstration of altered SBI following exogenous drug treatment. Notably, only inhibitory compounds enhanced goal-directed activity, linking glutamatergic attenuation to performance. Neurocomputational analysis revealed nuanced pharmacological responses during closed-loop stimulation, highlighting insights beyond spontaneous activity metrics.

## 1 Introduction

Predicting the clinical efficacy of a pharmacological intervention targeting a disease with *in vitro* microphysiological systems (MPS) requires viable assays that physiologically capture the key processes and functions of the tissue or organ it proposes to model. It is likely that no other biological system is as challenging to meaningfully model as neural systems. Neurons are highly dynamic cell types whose primary function is to send, receive and transform electrical signals throughout the brain and periphery. Current MPS models typically provide no stimulation to neural cells, with even state-of-the-art approaches using spontaneous activity or basic evoked activity as a functional outcome [1]. Yet, neural systems do not merely fire action potentials, but process structured information and alter both local and population-wide activity in meaningful ways. As a result, *in vitro* assays which examine the information processing capabilities of neurons (termed synthetic biological intelligence, or SBI) have recently been described (for a review of SBI and recent advances, see [2]). To process information effectively, neural systems must also be able to respond to a variety of cellular, environmental, and indeed pharmacological inputs with meaningful alterations in their activity [3–6]. Observing the results of such modulations would be critical for SBI assays to be useful as disease models or as drug discovery screening tools, yet an outstanding question is whether the learning seen in SBI assays is subject to pharmacological intervention.

In this study, we therefore assess for the first time whether function in the recently described MPS-based SBI assay known as *DishBrain* [7] can be modulated by the use of pharmacological compounds. We propose an SBI assay will provide the opportunity to assess not only the functional properties of a disease model, but also nuances that arise in response to pharmacological intervention. By creating a structured information landscape through a real-time closed-loop patterning of electrophysiological stimulation and recording, *in vitro* neural cultures can be embodied within a simulated game-world environment, in this case representing a simplified version of the game Pong (see Supplementary Methods 1 and Supplementary Figure 1 for details of the *DishBrain* closed-loop system, stimulation and feedback schemes). The observation of learning effects has been since replicated by another group, which supports the idea that exploring information processing in *in vitro* neurons is a viable direction for future research [8]. Previous work also demonstrated that cultures of cortical neurons within the *DishBrain* system displayed activity changes consistent with goal-directed learning, with significant increases in gameplay performance over time [7]. Moreover, these networks displayed population network activity metrics such as close to critical dynamics [9]. However, prior work has focused on *in vitro* neural cultures intended to model phenotypically “healthy” neural activity. Exploring the behavior of neural cell cultures designed to model disease characteristics would determine the viability of combining an MPS approach with SBI assays to test disease models and screen pharmacological interventions [10, 11].

Glutamatergic hyperactivity dysregulation is a state where excitatory forebrain neurons’ activity is high frequency, continuous and non-synchronized [12]. Post-mortem studies have associated irregular glutamate signaling with a number of neurological and psychiatric conditions [12]. A disease commonly associated with glutamatergic hyperactivity is epilepsy [13, 14]. Despite several disease-modifying therapies being available, approximately 30% of patients with epilepsy do not respond to these treatments [15]. As with many other neurological diseases, pre-clinical research to develop new ASMs has predominantly leveraged rodent models [16, 17]. Limitations associated with the use of these models have been identified and may, in conjunction with the complexity of the underlying condition and associated pathobiology, contribute to poor clinical trial outcomes [18]. At the same time, *in vitro* epileptiform cellular models using human neurons have been developed [13]. The ability to demonstrate compelling dose-response relationships in these *in vitro* models would be highly useful for drug discovery. Moreover, this could enable a personalized medicine approach: culturing cells from patients who display drug-resistant phenotypes; it would also reduce the need for animal testing.

Previous work has leveraged neurogenin 2 (NGN2)-reprogrammed neurons (cultured on an astrocyte layer) to create an *in vitro* model of epilepsy [19, 20]. By utilizing a robust forward programming platform for Optimized inducible Overexpression (OPTi-OX) of NGN2, human induced pluripotent stem cells (hiPSC) can reproducibly generate neuronal populations with glutamatergic hyperexcitability [21, 22]. However the extent that these glutamatergic hyperactive cultures show information processing changes and whether these cultures may exhibit changes in response to ASMs remains unknown. Here we aim to use these neuronal cultures that demonstrate glutamatergic hyperexcitability and explore the validity of the system as a simple *in vitro* model of epilepsy. By assessing the electrophysiological development of these cultures over time, including response to specific ASMs, we evaluated the suitability and utility of the *DishBrain* system for exploring disease modelling and drug screening. Finally, given the role of neurocomputational metrics such as neural criticality in predicting phenotypically normal behaviors *in vivo*, and the link these metrics have to epilepsy, we also assessed the population dynamics of these cultures during spontaneous activity and during gameplay in the *DishBrain* system [9, 23, 24].

Specifically, we hypothesized that NGN2-reprogrammed iNeuron cultures would demonstrate increased gameplay performance and phenotypically normalized electrophysiological population dynamics following the administration of existing anti-seizure drugs.

## 2 Materials and Methods

### iPSC Culture

A broad overview of the study methodology including cell culture can be seen in Figure 1 a). All cultures were maintained in a 37^◦^C, 5% CO_2_, 5% O_2_ humidified cell culture incubator (BINDER, Germany). NGN2 iPSCs were thawed on vitronectin (VTN, ThermoFisher, Australia)-coated T25 flasks (18 *µ*L VTN to 1.8 mL Dulbecco’s phosphate buffered saline, DPBS, containing Ca^2+^/Mg^2+^, washed with 4 mL DPBS without cations, -/-, prior to cell seeding) and maintained in 4 mL StemFlex iPSC media (ThermoFisher, Australia). Cells were passaged using EDTA (0.5 mM in DPBS -/-).

**Fig. 1:**
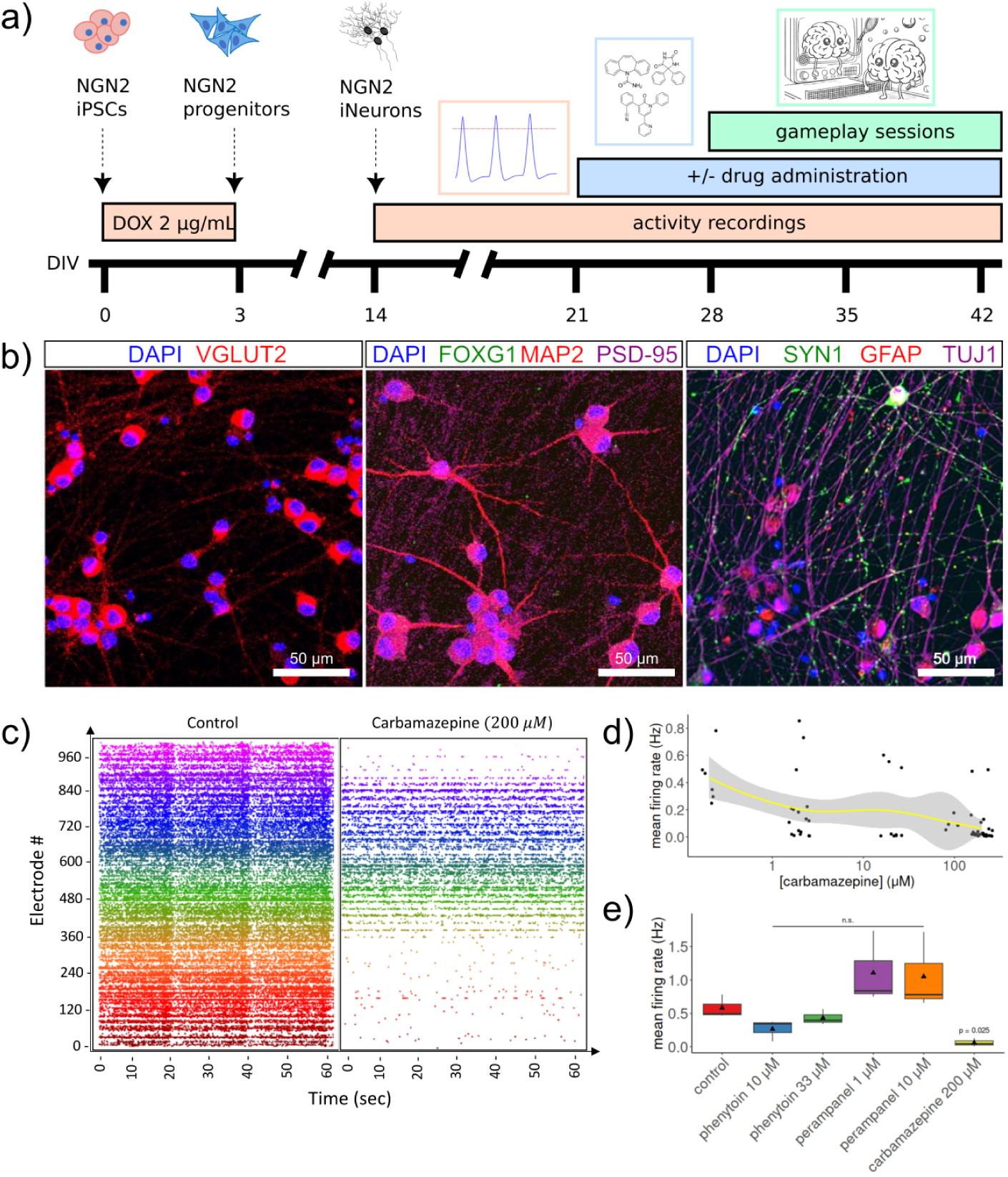
Overview of study design and basic NGN2 culture characterisation. a) Study overview. b) Immunocytochemistry of NGN2 cultures at day 38 showing brightfield, DAPI (blue), and VGLUT2 (SLC17A6; red) labelling. Immunocytochemistry in NGN2 cultures showing brightfield, DAPI (blue), FOXG1 (green), MAP2 (red), and PSD-95 (DLG4; magenta) labelling. Immunocytochemistry of NGN2 cultures at day 38 showing DAPI (blue), SYN1 (green), GFAP (red), and TUJ1 (magenta) labelling. Scale bar, 50 *µ*m. c) Raster plot showing culture firing activity prior to and after carbamazepine (200 *µ*M) administration at day 28. Each row represents one electrode (of 1024). Each point in the plot represents one detected spike. d) Relationship between mean firing rates and escalating doses of carbamazepine. Firing rates were calculated per 100 msec for this analysis. e) Analysis of mean firing rate with administration of different pharmacological compounds. Each group was analyzed by one-way ANOVA with post-hoc Bonferroni test compared to control. Box plots show interquartile range, with bars demonstrating 1.5X interquartile range, the line marks the median and the black triangle marks the mean.

### Neuronal Differentiation

To initiate differentiation, NGN2 iPSCs [22] were dissociated with Accutase (Merck, Australia) and seeded on Geltrex (ThermoFisher, Australia)-coated 24 well plates at a concentration of 80000 cell per cm^2^ in 1 mL StemFlex media with 5 *µ*M Y-27632. The next day (day 0), cells were washed with 1 mL DPBS -/- and media changed to Day 0-1 media (Table 1) containing doxycycline 2 *µ*g / mL (DOX, Merck, Australia). Cells were maintained in this media until day 2, when cells were washed with 1 mL DPBS -/- and media changed to Day ≥ 2 media (Table 1) containing DOX 2 *µ*g / mL. On day 3, cells were washed with DPBS -/-, dissociated with Accutase and plated on to Poly-D-Lysine-Geltrex coated MaxONE multi electrode array chips (Maxwell BioSystems, Switzerland) in Day ≥ 2 media (Table 1) containing DOX (2 *µ*g / mL) and 5 *µ*M Y-27632. 200000 NGN2 cells, and 50000 primary human cortical astrocytes (Sciencell Cat# 1800, USA; maintained previously according to manufacturer’s specifications) were mixed and plated on each MEA chip at this time. On days 4 and 5, full media changes in Day ≥ 2 media + DOX were performed. On day 7, media was changed to Day ≥ 2 media without DOX. Thereafter, half media changes were performed every second day.

**Table 1:**
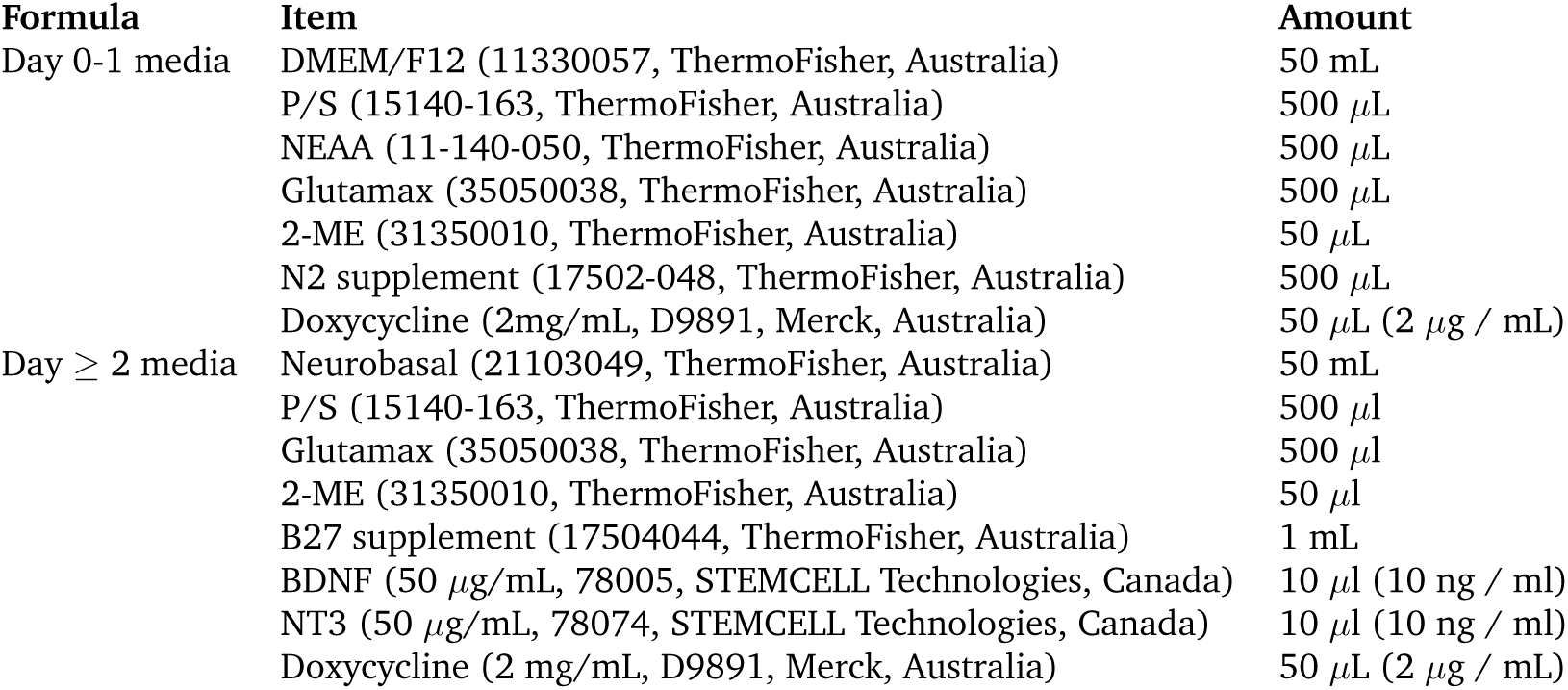
Cell culture media used in neural differentiation in the present study.

### Immunocytochemistry

From day 35 onwards, cultures grown in parallel on plastic tissue culture-treated 24-well plates were washed free of media (3 × 5 min in DPBS -/-) and fixed in 4% paraformaldehyde aqueous solution (Electron Microscopy Services, USA) for 15 min at room temperature (RT; 20°C). Wells were washed once with DPBS -/- and then permeabilized with 0.1% Triton X-100 (Merck, Australia) in DPBS -/- for 30 min at RT. Cells were then blocked with 1% normal goat serum in DPBS -/- for 30 min and incubated overnight at 4°C in 0.1% Triton X-100 in DPBS -/- with the primary antibodies shown in Table 2. The next day, after another 3 washes, secondary antibodies as shown in Table 2 were applied for 2 h at room temperature. Cells were again washed 3 times and the nucleic acid stain DAPI (ThermoFisher, Australia) was briefly added for 5 min prior to visualization to indicate the presence of cells. Cultures were visualized using a Nikon A1Plus Ti ZDrive confocal microscope (Nikon, Japan), and at least three images (2048 x 2048 pixels with 161 z-steps of 0.7 *µ*m) were recorded per well.

**Table 2:** Antibodies used in immunocytochemistry experiments.

| Group | Antibody | Usage |
| --- | --- | --- |
| Primary | Anti-VGLUT2 (ab79157, Abcam, USA) | 1:1000 |
|  | Anti-FOXG1 (ab196868, Abcam, USA) | 1:100 |
|  | Anti-MAP2 (ab5392, Abcam, USA) | 1:5000 |
|  | Anti-PSD-95 (ab192797, Abcam, USA) | 1:1000 |
|  | Anti-SYN1 (ab254349, Abcam, USA) | 1:500 |
|  | Anti-GFAP (ab4674, Abcam, USA) | 1:500 |
|  | Anti-TUJ1 (MAB1637, Merck, Australia) | 1:1000 |
| Secondary | Gt anti rabbit 488 (ab150077, Abcam, USA) | 1:1000 |
|  | Gt anti chicken 555 (ab159170, Abcam, USA) | 1:1000 |
|  | Gt anti mouse 647 (SAB4600183, Merck, Australia) | 1:1000 |
| Stain | DAPI 1 mg / mL (62248, ThermoFisher, Australia) | 1:10000 |

### Single Cell RNA Sequencing

Human iPSCs (D0) and iNeurons (days 7, 14, and 21) were washed once with 1X DPBS-/- before adding either Accutase (day 0) or Accutase containing 20 units / mL of papain, 5 mM MgCl_2_ and 5 mg / mL DNase I (Worthington, United Kingdom) (days 7-21). The cells were incubated at 37^◦^C for up to 5 min (day 0) or up to 45 min (days 7-21) before adding DMEM/F12 (ThermoFisher, United Kingdom) supplemented with 10% fetal bovine serum (Merck, United Kingdom). The cells were dissociated using a P1000 and collected in a 15 mL tube. After centrifugation, the cells were resuspended in 1X DPBS -/- containing 1% BSA (Merck, United Kingdom) and 10 *µ*M ROCK inhibitor (Y-27632, Tocris, United Kingdom) and collected in a 15 mL tube capped with a 40 *µ*m cell strainer. Following centrifugation, cells were washed three additional times. Single-cell suspensions were counted using an automated cell counter (Countess, Invitrogen, United Kingdom) and concentrations adjusted to 1000 cells / mL. Cells were then hash-tagged using total-seq anti-human hashtag antibody (BioLegend, United Kingdom) and kept on ice for further processing.

Single cell suspensions were processed by the Chromium Controller using Chromium Single Cell 3’ Reagent Kit v2 by 10x Genomics. Each sample was loaded for a target recovery of 8000 cells per sample. All the steps were performed according to the manufacturer’s specifications. All samples were sequenced using the Illumina HiSeq platform (Illumina, USA), with 50,000 reads on average per cell. scRNAseq samples were processed using the 10x Genomics CellRanger pipeline v3.0.1 using the GRCh38 genome assembly to produce the gene expression matrices. The filtered gene expression matrix was used without modifying any cell gating parameters.

Standard quality control was performed in R (v.4.3.1) using Seurat (v.4.4.0): Cells with mitochondrial RNA below 5% and with 200 to 10000 genes were included. Counts were scaled and mean expression levels at each timepoint were normalized to day 0 (hiPSCs) for visualization.

### Electrophysiology and *DishBrain* Pharmacology

MEA chip activity was assessed by performing activity recordings using the MaxWell Max-ONE system and software (Maxwell BioSystems, Switzerland) at regular timepoints during neuronal differentiation. Briefly, MEA chips were removed from the cell culture incubator and placed on one of three MaxONE readers in a non-humidified 37^◦^C 5% CO2 incubator and equilibrated for 10 min before 5 min of full recording. As reported previously [7] when chips obtained a culture-wide average activity of 0.7 Hz (generally between day 21 and day 35 of differentiation), chips were considered ready for *DishBrain* gameplay. On days where gameplay was performed, cells were removed from the cell culture incubator and an activity assay conducted as above to ensure minimum required activity for gameplay was present. If the chip passed, cells were treated with one of the following compounds (all using DMSO as a solvent, and diluted such that final DMSO concentration did not exceed 0.1% (v/v)): phenytoin 10-33 *µ*M, perampanel 1-10 *µ*M, carbamazepine 2-200 *µ*M, before being returned to the cell culture incubator. For vehicle control chips, DMSO was added at a final concentration of 0.1% (v/v). After 1 h, cells were returned to the reader for another activity assay, then closed-loop feedback gameplay was started as follows: Rest session (15 min), Active session (15 min), Rest session (15 min), Active session (15 min). Afterwards, chips had a full media change (1 mL) and were returned to the cell culture incubator. In Rest sessions, cell responses were recorded and contributed to gameplay, but cells were not stimulated with information encoding the gameplay world. In Active sessions, this information was also provided to the cells. (When refering to “gameplay” generally, we are talking about the *DishBrain* assay itself, and in the context of gameplay results, we are describing the culture’s performance in the Active session with respect to the Rest session, unless specifically noted in the text or figure. Further details on stimulation and configuration parameters during Rest and Active gameplay are presented in Supplementary Methods.) Chips were exposed to 60 min gameplay conditions a maximum of one time per day, and played for up to four consecutive days.

### Data Analysis

Activity assay recordings were analyzed to measure average spike firing rate and average spike amplitude. From gameplay experiments, hit/miss ratio and rally length were calculated and results summarized for each treatment (drug/dose) group. Summary data is presented as the mean hit/miss ratio or rally length, the standard error of the mean (SEM), and the count (n) of the treatment observations. Significance testing was performed to determine differences, if any, between treatment groups (Student’s t-test or one-way ANOVA with post-hoc Bonferroni multiple comparison test compared to DMSO vehicle control). A p value less than 0.05 was taken as the minimum level of significance. The majority of data analysis was performed in Python as described in section results. Initial electrophysiological analyses were performed using the RStudio (Posit Software, PBC) Build 463 IDE with R version 4.3.1 and the following packages: tidyverse version 2.0.0, ggsignif version 0.6.4.

## 3 Results

### NGN2 iNeuron cultures display glutamatergic and neuronal markers

To confirm the glutamatergic nature of the OPTi-OX NGN2 iNeurons, we began by examining common neuronal markers at the transcriptional and protein-levels. Immunocytochemistry of cultures on day 38 (a time when the cells were clearly active) revealed the presence of the neuronal markers MAP2 and TUJ1, the synaptic proteins SYN1 and PSD-95, as well as a high level of expression of VGLUT2 (SLC17A6), a marker of glutamatergic neurons (Figure 1, b, with individual channel images shown in Supplementary Figure 2, a, c, and d). With respect to VGLUT2, we segmented DAPI and VGLUT2 objects using FIJI imaging software and quantified the respective overlap (Supplementary Figure 2, b). We also confirmed the presence of GFAP, the glial fibrillary protein present in astrocytes (Supplementary Figure 2, d). We did not observe robust immunolabelling from FOXG1, a transcription factor required for ventral telencephalic development (Supplementary Figure 2, c). Single cell RNA-Seq experiments further confirmed the upregulation of neuronal, synaptic, and glutamatergic genes, as well as the downregulation of pluripotency genes, from day 0 to day 21 of differentiation. SLC17A7 (VGLUT1) and NGN2 expression peaked at day 7 before returning to baseline, and FOXG1 expression did not appear to change during the timepoints measured (Supplementary Figure 3).

### NGN2 iNeuron cultures display electrophysiological hyperactivity and pharmacological interventions showcase compound-specific responses

Having confirmed the glutamatergic identity of the neurons in culture, we turned our attention to characterizing their basic functional properties. Cultures developed rapidly over the course of two weeks, from day 7 to day 21, and displayed robust burst and network activity in all channels (Figure 1, c, shows control activity recorded at day 28). We then measured their activity after the application of carbamazepine, phenytoin, and perampanel. Carbamazepine had a marked effect on reducing mean firing rate after 1 h treatment (Figure 1, c), and we observed a dose-response relationship between increasing carbamazepine dose and reduced mean firing rate (Figure 1, d). At 200 *µ*M, the reduction in firing activity brought about by carbamazepine treatment was significant (Figure 1, e; *p <* 0.05), however phenytoin and perampanel did not significantly affect the firing rate of cultures at either dose tested.

We next investigated the electrophysiological activity of NGN2 cultures during gameplay, and analyzed the mean firing rate calculated per second, the average variance of the extracted firing rates and the average interspike intervals (ISI) among recorded channels on every culture. The distribution of these metrics is depicted in Figure 2 (a, c, e), showcasing comparisons between gameplay and rest conditions under control, as well as across various pharmacological interventions. The plots show an estimated probability density function using Kernel Density Estimation (KDE) with Gaussian smoothing. The negative values in the KDE plots arise from Gaussian smoothing, where the density estimate extends beyond the zero boundary due to data near zero. Statistical comparisons are shown in Figure 2 (b, d, f). Additionally, we explored the same electrophysiological metrics across different doses and compared these to the control group as seen in Supplementary Figure 4 (a, b, c). iNeuron cultures also displayed regular synchronised network activity (Supplementary Figure 5) at rest and after treatment with the pro-convulsant 4-aminopyridine (4-AP).

**Fig. 2:**
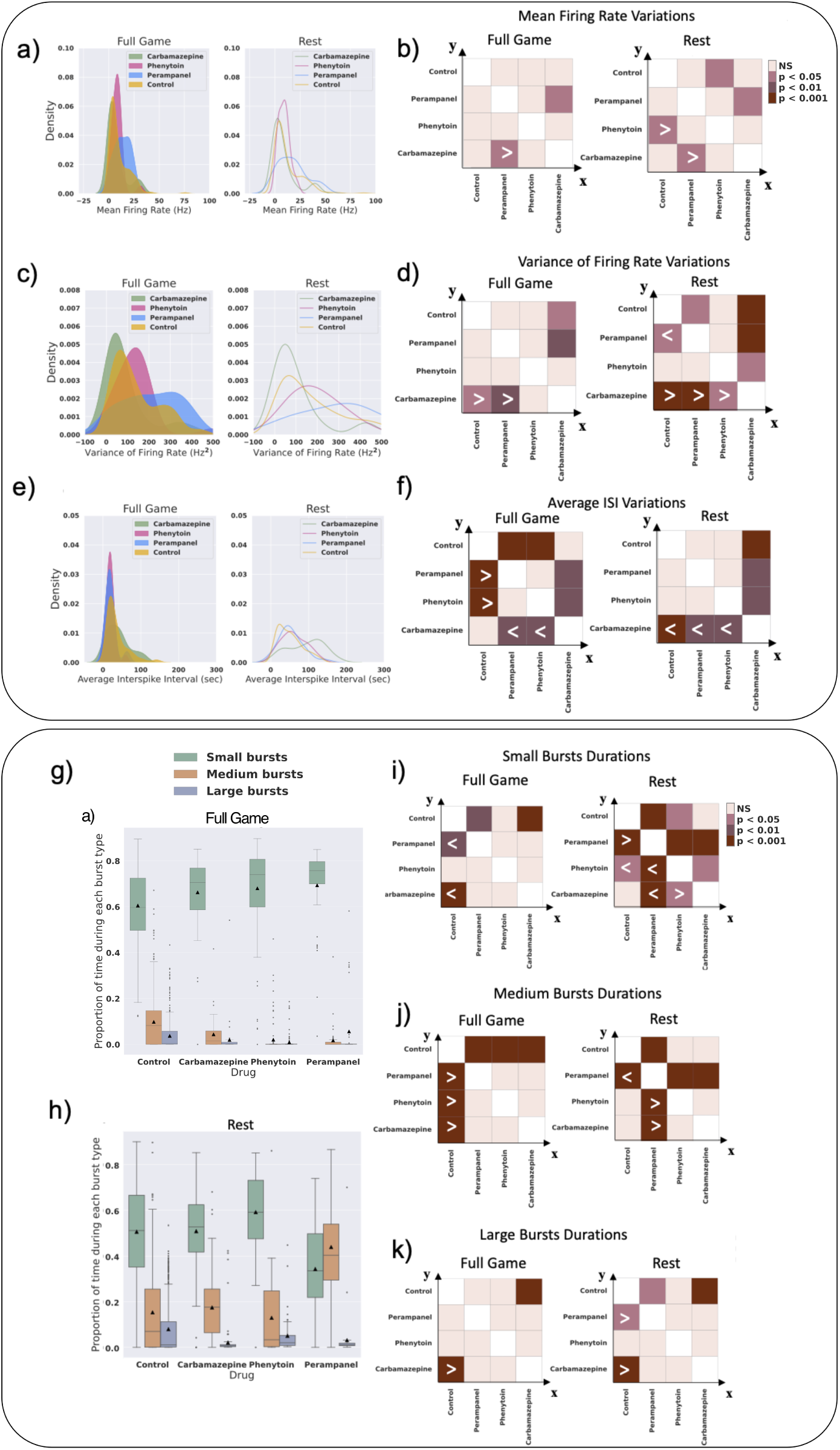
Neural firing and burst dynamics under control and pharmacological intervention conditions. a, c, e) Estimated probability density function plots using KDE with Gaussian smoothing to compare Mean Firing Rate, Variance of Firing Rate, and Average Interspike Interval (ISI) in both rest and full game sessions. The negative values in the KDE plots arise from Gaussian smoothing, where the density estimate extends beyond the zero boundary due to data near zero. The sample sizes (number of recording sessions) are n = 42, n = 42 (Carbamazepine: rest, full game), n = 24, n = 24 (Phenytoin: rest, full game), n = 24, n = 23 (Perampanel: rest, full game), n = 354, n = 400 (Control: rest, full game). b, d, f) Post-hoc Games-Howell test between all the rest and all the full game groups. *>*: *x > y* by significance level and *<*: *x < y* by significance level. g, h) Boxplots illustrating the average total proportion of time spent during the occurrence of each burst type (small, medium, and large) under control and pharmacological intervention conditions. sessions. The sample sizes are n = 354, n = 400 (Control: rest, full game), n = 42, n = 42 (Carbamazepine: rest, full game), n = 24, n = 24 (Phenytoin: rest, full game), and n = 24, n = 23 (Perampanel: rest, full game). Box plots show interquartile range, with bars demonstrating 1.5X interquartile range, the line marks the median and the black triangle marks the mean. Error bands = 1 SE. i, j, k) The significance heatmaps depict the post-hoc Games-Howell test results comparing control and different pharmacological intervention conditions during rest and the full game. On the axes, “*>*” indicates that the group on the *x*-axis is greater than the group on the *y*-axis at the given significance level, and “*<*” indicates that *x < y* at the given significance level.

Drawing inspiration from the burst classification methods outlined in Wagenaar et al., [25], we extracted quantitative details from the bursting patterns of the recordings from each *in vitro* culture. In order to determine whether the glutamatergic hyperactivity in our cultures was linked to synchronized burst activity, we examined the amount of time each ASM treatment group was in a burst state (Figure 2, g and h). Significant changes in small, medium, and large burst states respectively are shown in (Figure 2, i, j, and k).

### Analysis of gameplay performance under control and pharmacological intervention conditions

To evaluate the game performance of cultures exposed to various pharmacological compounds, three distinct metrics were employed: Average Rally Length (representing the average number of accurate hits per rally), Hit/Miss Ratio (indicating the ratio of accurate hits to missed balls), and Aces to Game Ratio (illustrating the proportion of games where the ball is immediately missed after the initial serve, known as an ace). The estimated probability density functions of these game performance metrics using KDE are visualized across all groups. (Figure 3, a, c, e), and a statistical significance table is presented for comparing the mean values of different groups (Figure 3, b, d, f). Subsequently, the same metrics are computed for various doses of each administrated drug (Supplementary Figure 6, a, b, c). We next examined the effect of duration of gameplay on our analysis. Figure 3 (g, h, i) represents the percentage of changes in average rally length compared to rest under control and all the pharmacological interventions when studying the first and second half of each 15-min-long recording. The statistical significance table is presented for comparing the mean values of different groups in Figure 3 (j, k, l). The high dose of carbamazepine (200 *µ*M) was the only group showing a significant improvement in time in terms of the average rally length compared to rest.

**Fig. 3:**
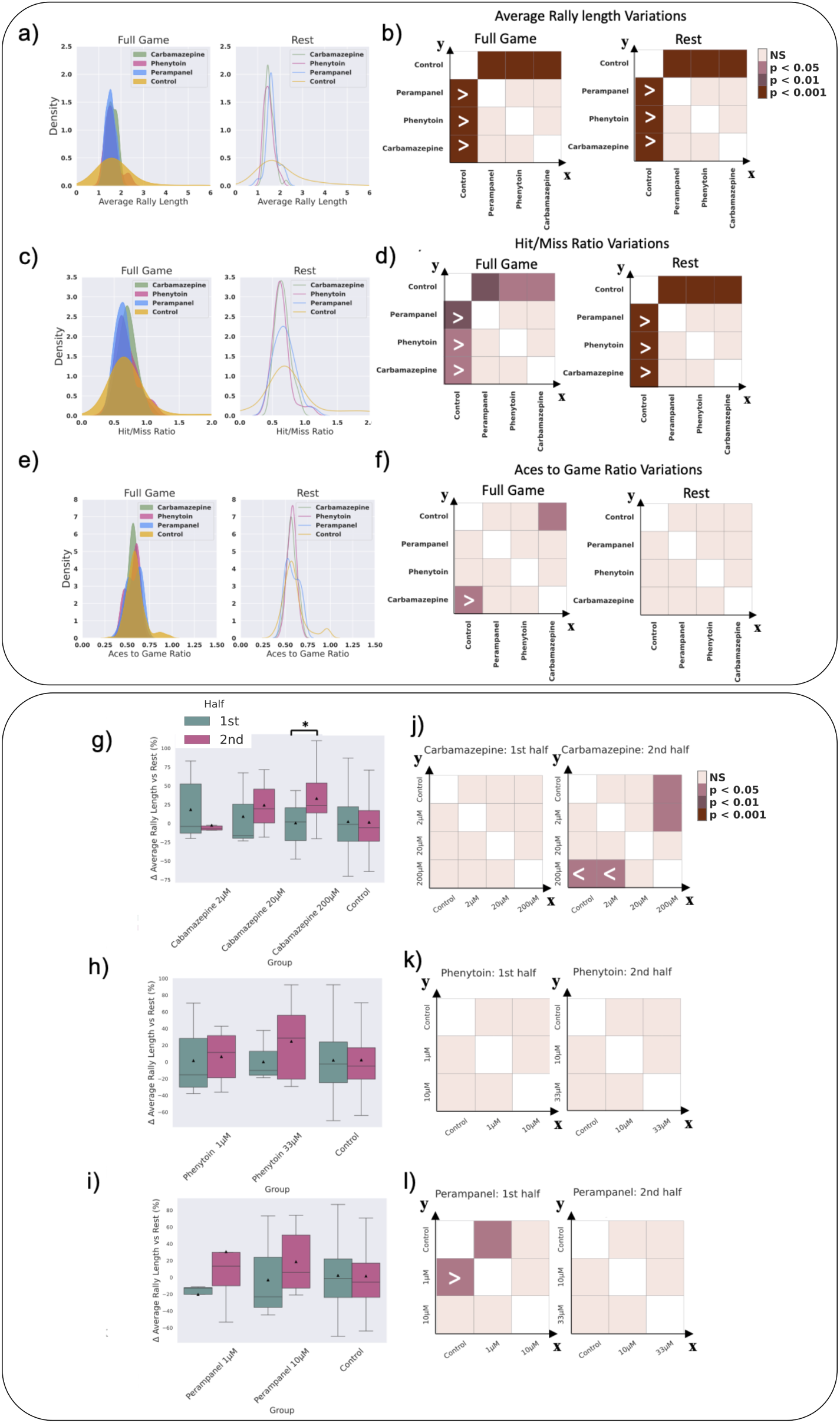
Game performance analysis under control and pharmacological intervention conditions. a, c, e) Comparison of Average Rally Length, Hit/Miss Ratio, and Aces to Game Ratio in both rest and full game sessions. Kernel Density Estimation plots are used, applying Gaussian smoothing to estimate the probability density function of the data. The sample sizes are n = 42, n = 42 (Carbamazepine: rest, full game), n = 24, n = 24 (Phenytoin: rest, full game), n = 24, n = 23 (Perampanel: rest, full game), n = 354, n = 400 (Control: rest, full game). b, d, f) The significance heatmaps depict the post-hoc Games-Howell test results comparing control and different pharmacological intervention conditions during rest and the full game. On the axes, “*>*” indicates that the group on the *x*-axis is greater than the group on the *y*-axis at the given significance level, and “*<*” indicates that *x < y* at the given significance level. g, h, i) Comparison of Average Rally Length changes in full game sessions vs rest when comparing the first and second half of each recording. * indicates *p <* 0.05. j, k, l) The significance heatmaps depict the post-hoc Games-Howell test results comparing control as well as various doses of each pharmacological intervention condition during the first and second half of full game sessions. On the axes, “*>*” indicates that the group on the *x*-axis is greater than the group on the *y*-axis at the given significance level, and “*<*” indicates that *x < y* at the given significance level.

### Bursting patterns of activity under control and pharmacological intervention conditions

We further explored bursting patterns of activity under both control and pharmacological intervention conditions as described above. In certain recordings, bursts displayed highly consistent sizes, while others exhibited a broad range of burst sizes.

We proceeded to tally the number of small, medium, and large bursts in all recordings. Furthermore, we normalized the counts of various burst sizes in all subsequent sessions to the count of those burst sizes in the initial rest session recording. The average duration of burst activity in recordings at rest and during gameplay is shown in Supplementary Figure 7. Figure 4 (a, b, c) shows the number of extracted small, medium, and large bursts from full game and rest sessions under control and all the pharmacological interventions. Post-hoc statistical significance for the comparisons between groups are visually represented in Figure 4 (d, e, f). The same metrics are then illustrated for various administrated doses of each drug compared to the control group in Figure 4 (g-o). The count of small bursts dropped significantly both during rest and gameplay with administration of all compounds. The count of large bursts also significant decreased in both carbamazepine and phenytoin groups during gameplay relative to control and significantly decreased in the phenytoin and perampanel-treated groups during rest compared to the control group. Figure 4 (p) shows the Pearson correlation values between each of the burst pattern metrics and different electrophysiological and game performance metrics including the metrics introduced previously. The corresponding p-values are illustrated in Figure 4 (q). Interestingly, the Mean Firing Rate appears to be negatively and significantly correlated with the number of small and large bursts in the cultures.

**Fig. 4:**
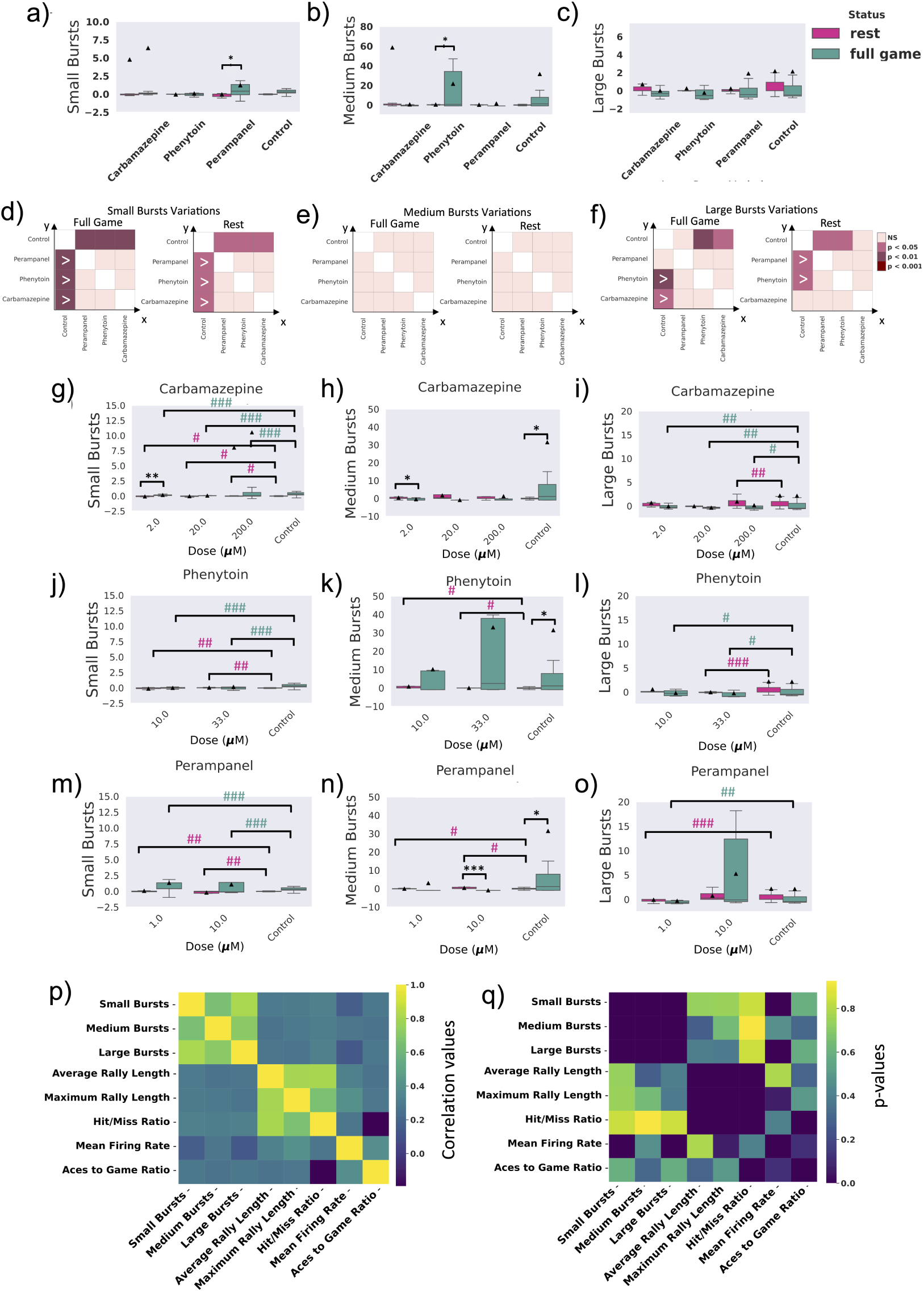
Burst pattern analysis under control and pharmacological intervention conditions. a, b, c) The normalized number of small, medium, and large bursts for rest and full game sessions of all control and drug treated cultures. The first rest session recording of each experiment is used for normalization. The sample sizes are n = 35, n = 35 (Carbamazepine: rest, full game), n = 20, n = 20 (Phenytoin: rest, full game), n = 20, n = 20 (Perampanel: rest, full game), n = 97, n = 95 (Control: rest, full game) from left to right. One-way ANOVA test. d, e, f) The significance heatmaps depict the post-hoc Games-Howell test results comparing control and different pharmacological intervention conditions during rest and the full game. On the axes, “*>*” indicates that *x > y* at the given significance level, and “*<*” indicates that *x < y* at the given significance level. g, h, i, j, k, l, m, n, o) Comparison of the same burst pattern metrics between different administrated doses of each drug. One-way ANOVA test was conducted within each group, and post-hoc Tukey’s test was conducted between all the rest and all the full game groups. # or * indicate *p <* 0.05, ## or ** indicate *p <* 0.01, ### or *** indicate *p <* 0.001. p, q) Pearson correlation test between the burst pattern and the game performance metrics.

### Critical dynamics of cultures under control and pharmacological intervention conditions

Dynamical systems undergo transitions between ordered and disordered states, and the concept of “criticality” emerges when the system resides at the boundary between these states—where the input is neither strongly damped nor excessively amplified [26]. Failures in adaptive criticality may contribute to impairments in brain function, such as those observed in dementia or epilepsy [27]. Building on insights from recent literature [9, 28], an avalanche analysis was conducted to examine the network in relation to its proximity to criticality.

The start and end of an avalanche are determined by crossing a threshold of network activity, with spikes from any or all neurons within a specified region capable of triggering an avalanche. The scale-free dynamics of detected neuronal avalanches, along with metrics such as the Deviation from Criticality Coefficient (DCC), Branching Ratio (BR), and Shape Collapse error (SC error), were assessed to gauge whether the recordings were poised near criticality [9]. A lower DCC and SC error, along with a BR closer to 1, serve as indicators of a state near criticality. Figure 5 (a, b, c) show the number of extracted DCC, BR, and SC error from full game and rest sessions under control and all the pharmacological interventions. Post-hoc statistical significance for the comparisons between groups are visually represented in Figure 5 (d, e, f). The same metrics are then illustrated for various administrated doses of each drug compared to the control group in Figure 5 (g-o). The results demonstrate that criticality metrics are greatly disrupted both under control and pharmacological intervention in both game states. Some small variations are observed such as decreased DCC and increased BR values in low doses of carbamazepine (2 *µ*M) or decreased SC error in higher doses (200 *µ*M) compared to control during gameplay. The BR also display a significant increase when comparing gameplay to rest state in 200 *µ*M carbamazepine. Overall, there is limited expression of the various criticality metrics in these cultures whose neurons are nearly all glutamatergic. This is supported by previous literature which showed that critical state dynamics arise from a balance of excitatory and inhibitory neuronal networks [26, 29].

**Fig. 5:**
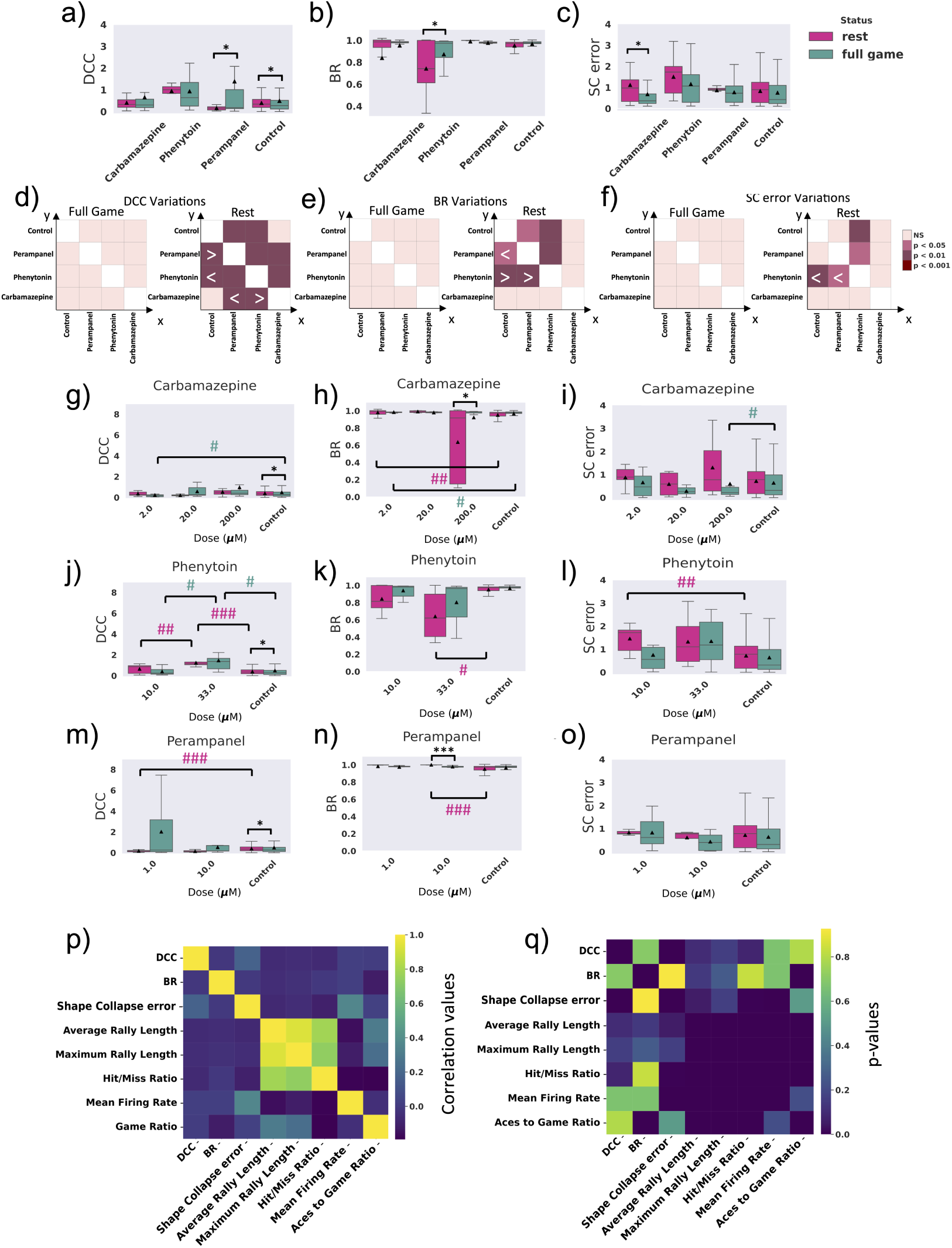
Criticality analysis under control and pharmacological intervention conditions. a, b, c) The Deviation from Criticality Coefficient (DCC), Branching Ratio (BR), and Shape Collapse error (SC error) for rest and full game sessions of all control and drug treated cultures. The sample sizes are n = 24, n = 36 (Carbamazepine: rest, full game), n = 20, n = 24 (Phenytoin: rest, full game), n = 12, n = 19 (Perampanel: rest, full game), n = 288, n = 369 (Control: rest, full game) from left to right. One-way ANOVA test. d, e, f) The significance heatmaps depict the post-hoc Games-Howell test results comparing control and different pharmacological intervention conditions during rest and the full game. On the axes, “*>*” indicates that *x > y* at the given significance level, and “*<*” indicates that *x < y* at the given significance level. g, h, i, j, k, l, m, n, o) Comparison of the same criticality metrics between different administrated doses of each drug. One-way ANOVA test withing each group and post-hoc Tukey’s test between all the rest and all the full game groups. # or * indicate *p <* 0.05, ## or ** indicate *p <* 0.01, ### or *** indicate *p <* 0.001. p, q, r, s) Pearson correlation test between the criticality and the game performance metrics. p, r) show the correlation values.

Figure 5 (p) shows the Pearson correlation values between each of the criticality metrics and different electrophysiological and game performance metrics including the metrics introduced previously. The corresponding p-values are illustrated in Figure 5 (q). Culture game performance measured in Hit/Miss Ratio is negatively and significantly correlated with SC error while Mean Firing Rate appears to have a positive and significant correlation with this criticality metric.

### Functional connectivity of cultures under control and pharmacological intervention conditions

In order to determine how ASM administration affected underlying network activity at rest and during gameplay, we used neuronal spiking activity to construct functional connectivity networks for all recordings. Due to the extensive duration of the recordings at a 20 kHz sampling frequency, the resulting time series data was substantial. Following the approach outlined in Khajehnajad [30], dimensionality reduction techniques, particularly t-SNE [31], were employed to enhance computational efficiency and improve data interpretability. The 3-dimensional representations obtained using t-SNE for both rest and gameplay recordings were utilized to investigate latent network structures. To evaluate the effectiveness of these low-dimensional representations, each rest and gameplay recording session was first divided into two equal time intervals prior to applying dimensionality reduction. Figure 6 (a, c, e, g) shows the results after color-labeling the first and second halves of the recording sessions for sample cultures under both gameplay and rest conditions. The visualization shows that while the two halves of gameplay sessions are distinguishable using t-SNE, the distinction is less evident during rest sessions. Functional connectivity network matrices were then constructed using zero-lag Pearson correlations for each gameplay or rest session recording, with the 1024 channels as nodes and weighted edges representing functional connectivity. Only edges with Pearson correlation absolute values above 0.7 were retained.

**Fig. 6:**
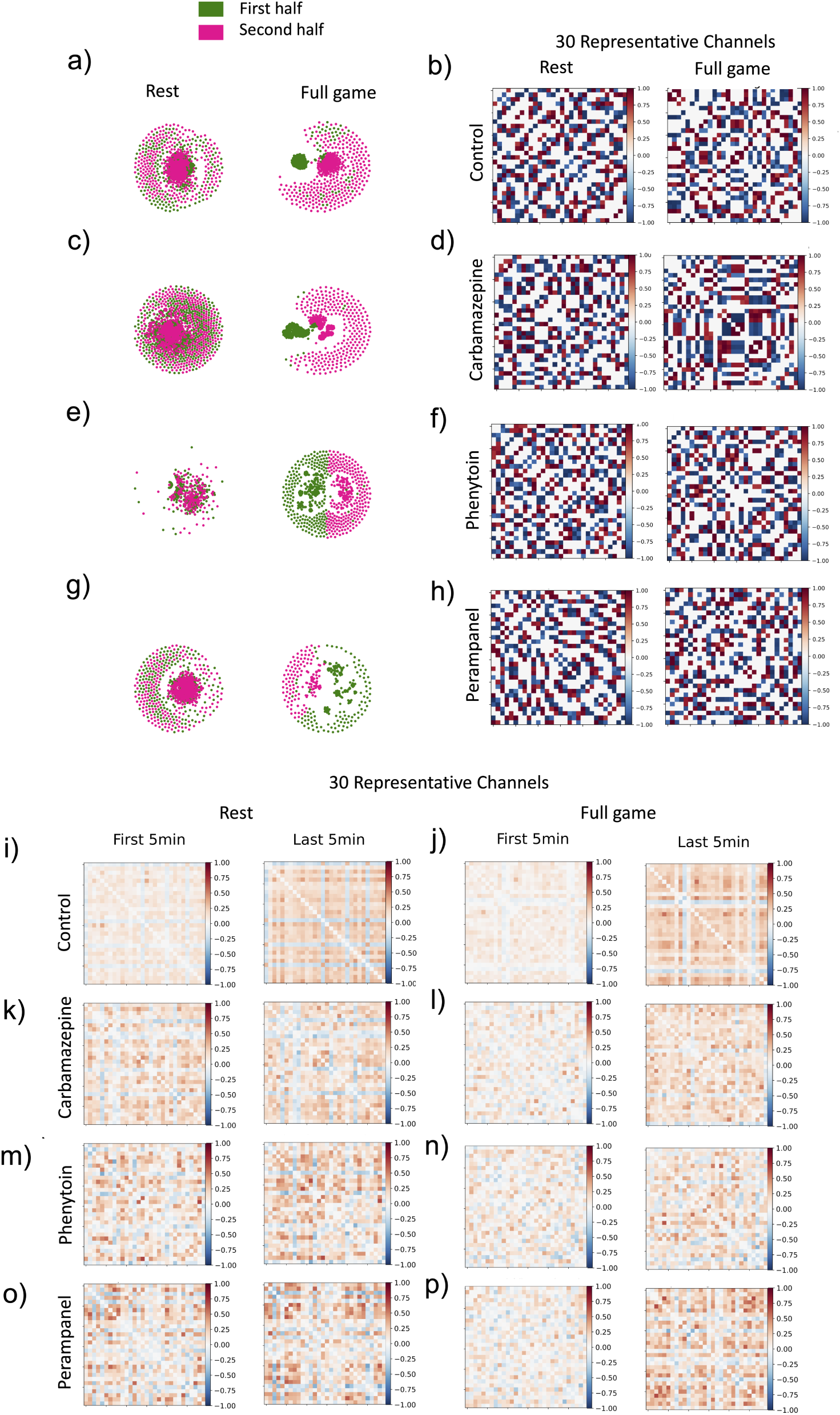
Functional connectivity networks of rest and full game sessions under control and pharmacological interventions. a, c, e, g) Low-dimensional representation of samples of rest and full game sessions using t-SNE under control and pharmacological intervention conditions (Carbamazepine, Phenytoin, and Perampanel, from top to bottom, respectively). The green and purple dots are the channel representations in the embedding space in the first and second halves of the recordings, respectively. b, d, f, h) Heatmaps illustrating the average edge weights between the 30 representative channels in the Pearson Correlation network constructed from all rest and all full game recordings under control and pharmacological intervention conditions. The sample sizes are n = 42, n = 43 (Carbamazepine: rest, full game), n = 24, n = 24 (Phenytoin: rest, full game), n = 24, n = 24 (Perampanel: rest, full game), n = 354, n = 400 (Control: rest, full game). One-way ANOVA test. i-p) Changes in the functional connectivity networks of rest and full game sessions under control and pharmacological interventions. Heatmaps illustrate the change in average edge weights between the 30 representative channels in the Pearson Correlation network constructed from the first and last 5 minutes of all rest and all full game recordings under control and pharmacological intervention conditions.

Considering that only a fraction of neurons fire at any given time within complex neural networks, the same method introduced in [30] was utilized to reduce computational complexity while preserving network dynamical properties. This involved identifying a subset of recorded channels that likely monitored neuronal populations specifically attuned to the ongoing task. Tucker decomposition via higher-order orthogonal iteration on tensor data from all lower-dimensional representations of the recordings was employed to identify a consistent subset of channels across all neuronal cultures. K-medoid clustering was then applied to partition the data into 30 clusters, and the corresponding ‘medoids’ were extracted as the mutual representative channels amongst all cultures. Finally, another network matrix using Pearson correlation was built with these 30 channels as nodes. The resulting functional connectivity networks were derived separately for the first and last 5 minutes of every recording.

Figure 6 (b, d, f, h) illustrate the heatmap of correlation values (i.e. edge weights) in the constructed functional connectivity networks for the smaller network (30 representative channels) and during rest and gameplay sessions. Heat maps of the full network (1024 channels) are found in Supplementary Figure 8. Next, to evaluate the dynamics of the functional connectivity networks in time, we divided each of the rest or gameplay sessions into 5-minute intervals and visualized the connectivity networks for the first and last 5-minutes of each group.

As a result of all pharmacological interventions, these functional connectivity networks using the low-dimensional representations of neural activity during rest and gameplay reveal distinct patterns evolving over time specifically during gameplay. This contrasts with the absence of such patterns during the spontaneous activity observed at rest as seen in Figure 6 (i-p). This difference is evident when considering both the full network (1024 channels; Figure 9) and the smaller network (30 representative channels; Figure 6 (i-p)).

We then studied various macroscopic metrics of the connectivity network for the control group and all the different doses of the pharmacological interventions. The measured metrics encompassed: 1. Average Weight: This metric illustrates the mean value of Pearson Correlations calculated between pairs of spiking time series in each culture. 2. Modularity Index: This index quantifies the extent to which a network can be partitioned into distinct and internally cohesive groups or communities. It evaluates how nodes form clusters that are more densely connected within themselves than with nodes outside the cluster. 3. Clustering Coefficient: This measure quantifies the tendency of nodes to form clusters or highly connected groups, providing insights into the local connectivity of the network by assessing the likelihood that neighbors of a particular node are connected to each other. Figure 7 (a-l) represents these derived network metrics for the gameplay and rest sessions under pharmacological interventions conditions, as compared to control conditions.

**Fig. 7:**
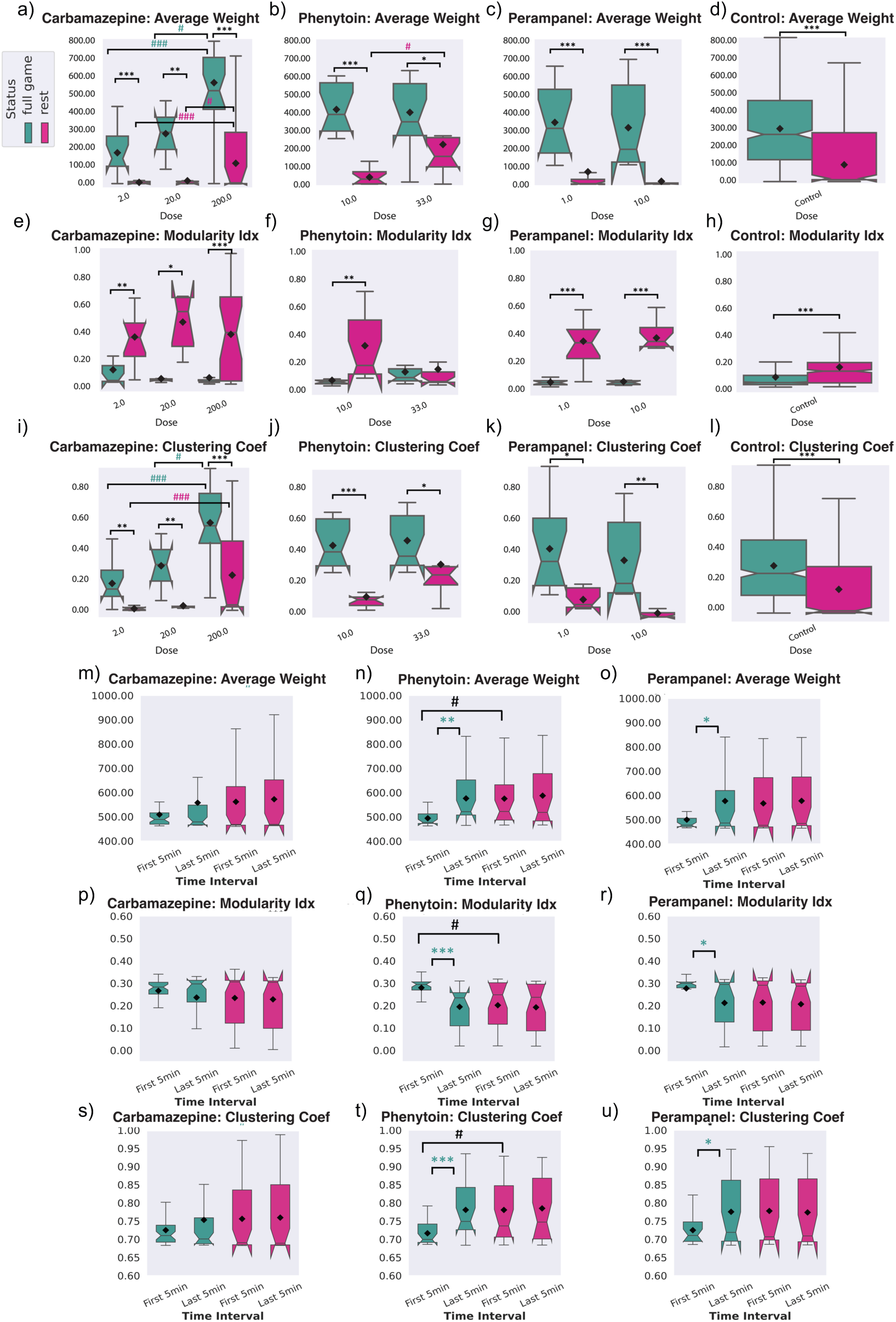
Network summary statistics using all of the recorded channels under control and different drug treatments. Comparing gameplay and rest recordings: a, b, c, d) Average weight, e, f, g, h) Modularity Index, and i, j, k, l) Clustering coefficient values for gameplay and rest recordings under control and all pharmacological interventions using the entire recording length from all 1024 recorded channels. One-way ANOVA test withing each group and post-hoc Tukey’s test between all the rest and all the full game groups. # or * indicate *p <* 0.05, # or ** indicate *p <* 0.01, ### or *** indicate *p <* 0.001. Network summary statistics between the first and last 5 minutes of full game and rest recordings using all of the recorded channels under different drug treatments. Comparing the first and last 5 minutes of full game: m, n, o) Average weight, p, q, r) Modularity Index, and s, t, u) Clustering coefficient values for the first and last 5 minutes of recordings under all pharmacological intervention using all 1024 recorded channels. One-way ANOVA test withing each group and post-hoc Tukey’s test between all the rest and all the full game groups. # or * indicate *p <* 0.05, ## or ** indicate *p <* 0.01, ### or *** indicate *p <* 0.001.

We then studied the changes in the introduced macroscopic network metrics from pharmacological interventions when comparing the first and last 5 minutes of recordings (see Figure 7 (m-u)). We utilize the first and last 5-minute time intervals, for both the full network and the smaller network. Supplementary Figure 10 represents the evolution of the macroscopic network metrics over time under the various pharmacological interventions using the full network. Supplementary Figure 11 studies the same network statistics for various doses of administrated drugs. Carbamazepine appears to show a significant change in network dynamics only when administered at its high dose (200 *µ*M). Increasing average weight and clustering coefficient as well as decreasing modularity index are observed during gameplay but not in rest for all of the administrated drugs. While these differences in gameplay are more significant for phenytoin and perampanel, specifically using the full network in Supplementary Figure 9, they rapidly lose the changes which arise during the initial embodiment in the gameplay environment —such as those observed in the first 5 minutes—and return to baseline levels observed during rest. This returning to baseline effect is not observed after carbamazepine administration.

## 4 Discussion

By using an *in vitro* iPSC neuron model characterized by glutamatergic hyperactivity, we intended to explore the behavior of these cells in a structured information landscape and the impact of pharmacological interventions intended to suppress electrophysiological hyperactivity. It was proposed that modelling dynamic environments through closed-loop electrophysiological stimulation and recording, would provide an enhanced MPS-based method to embody simple neural cultures in a simulated game-world and may offer additional information over existing drug-response assays traditionally used in MPS. The first step in this experiment was to establish a suitable disease model. Using the OPTi-OX system [22], NGN2-iNeurons were generated, which displayed elevated levels of glutamatergic markers. These NGN2-iNeurons demonstrated heightened spontaneous spiking, greatly exceeding that of primary cortical rodent cultures and dorsal forebrain cultures intended to model healthy activity [7]. It is also interesting to note that the spiking activity of these OPTi-OX inducible lines far exceeds the activity of virus-generated NGN2 cultures previously tested in the *DishBrain* system, which did show evidence of statistically significant learning over time without pharmacological intervention [7]. These results may reflect a greater consistency of NGN2 overexpression with the OPTi-OX system relative to viral methods, or instead that the OPTi-OX system results in similar transgene expression but with more hyperactivity in the resulting cultures; to conclusively determine this further work is required. Either way, these data suggest that OPTi-OX NGN2 iNeurons can produce a robust model of glutamatergic hyperactivity [13, 14, 19].

The evidence that these OPTi-OX NGN2 cultures were suitable as an epilepsy model was more nuanced. The NGN2 cultures displayed increased activity coupled with a limited inhibitory network. Some evidence of recurrent epileptiform activity, with large network bursts comparable to seizures and suppressed upon ASM administration, was also observed, but the complexity of this activity was limited due to the simple networks within these cultures. Further support for this model’s functionality was observed in the compound-specific electrophysiological modulation achieved when various ASMs were introduced to the cultures. Preliminary pharmacological treatments of these cultures showed a reduction in culture activity and changes in bursting patterns after an hour of exposure to ASMs. Interestingly, the magnitude of bursting pattern changes, particularly for large bursts, differed between ASMs. As expected, we observed concentration-response relationships between the dose of drugs administered, and mean culture firing rate, indicating the presence of the drug’s respective receptor targets, and the sensitivity of the glutamatergic hyperexcitable phenotype to pharmacological manipulation. The mechanism of action of all three compounds tested is established: perampanel binds non-competitively and selectively to AMPA-type glutamate receptors [32], and both phenytoin and carbamazepine block voltage gated sodium channels on the surface of neuronal cell membranes [33–35]. The effect of ASM administration is a reduction in aberrant neuronal excitability associated with depolarization and action potentials leading to a decreased likelihood of seizures. One limitation of our study is that, while the neural systems were supplemented with astrocytes, the model lacks other supporting cell types including interneurons and an appropriate balance of inhibitory cells which are likely important to capture more nuances expressed in epilepsy clinically. Despite this limitation, the reduction in firing rate and changes in burst state we measured in this study is consistent with the mechanism of action these compounds display clinically, and led us to question if the acute drug effect would translate to modulation of gameplay performance in the full *DishBrain* gameplay assay.

When examining the overall gameplay performance characteristics of the neural cultures, significantly more variance was observed in control cultures without pharmacological intervention. Despite this variation, mean culture performance, after treatment with carbamazepine, was significantly improved (more hits, longer rallies, fewer aces) when compared to control and other treatment groups. Similarly, only the 200 *µ*M carbamazepine group showed overall significantly higher performance on average rally length in gameplay compared to rest. This result becomes more pronounced when looking at learning over time instead of just between groups overall, consistent with previous work in this area [7, 36]. Despite these statistically significant differences, the effect sizes are notably smaller than in previous work. Of note, no statistically significant learning effects were observed over time in untreated cultures. Given the substantially different electrophysiological profile of the OPTi-OX NGN2-iNeurons tested in this work relative to the virally induced NGN2 cells tested previously, this may reflect the possibility that hyperactivity is not conducive to information processing and learning. This is supported by the observation that these small but significant learning effects occurred after the loss of the majority of activity in the presence of 200 *µ*M carbamazepine. If so, this would provide some support for the use of these cells as a disease model, in-line with previous work [19]).

Moreover, extending beyond whether neural cultures showcased any overt evidence of improved gameplay performance, it is also interesting to compare and contrast different electrophysiological phenomena while cells are embodied in the structured information landscape vs engaged only in spontaneous activity. For example, looking at different sizes of network burst activity, an increased dosage of phenytoin appears to significantly increase medium bursts during gameplay vs controls and the lower dose, but not during rest; perampanel has a similar effect on small bursts. These effects appear reasonably consistent across different dosages and likely reflect the nuances of the various compounds’ actions on neural function both with and without structured stimulation. Without the ability to embody neural cultures in these closed-loop systems, it would not be possible to extract this data, providing further evidence as to the benefit of augmenting neural MPS systems with this approach for future studies. Examining simple pairwise correlations did not reveal a direct and obvious link between these bursting patterns and gameplay characteristics. However, future work focused on explaining these mechanisms could aim to build more nuanced computational models to understand whether the existence of any moderated or mediated pathways do exist. Similarly, previous work established that when neural systems intended to model healthy activity were placed in a closed-loop structured information landscape they would show significantly closer to critical dynamics compared to when engaged in spontaneous activity alone [9]. Given that criticality has been identified as a clinically relevant metric and is specifically associated with epileptiform activity, it was a meaningful metric to consider in this work as well [23, 24]. Consistent with this previous literature, criticality was found to be greatly disrupted in these cultures, even though some small evidence of variations due to pharmacological intervention and game state was found. The limited expression of the various criticality metrics in these cultures whose neurons are nearly all glutamatergic also supports previous literature which showed that critical state dynamics arise from a balance of excitatory and inhibitory neuronal networks [29, 37]. Thus, the lack of excitatory-inhibitory balance in this work may explain the limited observations of critical dynamics and learning effects, relative to previously reported levels [9]. Despite this, significant correlations were still observed between some criticality and performance metrics. While this was not replicated across all measures, it provides support for further consideration of critical dynamics as a marker of pre-clinical pharmacological intervention.

Finally, we also explored the functional connectivity within neural cultures both unstimulated and when embodied in the gameplay environment. Previous research has found that functional connectivity is linked to key behavioral changes within *in vitro* cultures when embodied in structured information landscapes [8, 30]. This work is replicated here using low-dimensional representations of neural activity. Clear differences can be observed during gameplay, which are not seen during the purely spontaneous activity displayed during rest. This represents a degree of reorganization in neural cultures, even if such organization does not extend towards robust metrics of improved gameplay performance. When examining the average edge weights, either looking at all channels or the condensed 30 representative channels, it is apparent that there exists far more structure in the changes occurring in the cultures during the gameplay compared to rest, where minimal structured changes appear. Nonetheless, while looking at the network summary statistics between different pharmacological interventions during gameplay and rest a more nuanced picture emerges. In particular, it appears that administration of either phenyotin and perampanel results in the average weight, modularity index, and clustering coefficient metrics rapidly losing the changes which arise during the initial embodiment in the gameplay closed-loop environment, and returning to baseline levels observed during rest. In contrast, carbamazepine administration results in more stable networks that do not return to resting baseline. The use of more controllable MPS paradigms – ideally allowing for prolonged testing times – will be required to interpret these results and forms an important direction for future research. Providing mechanistic interpretations of nuanced changes observed in these dynamic systems exceeds the scope of this specific research. Despite this limitation, these results do showcase the utility of conducting pharmacological assays in neural cultures when assessing SBI characteristics in a closed-looped structured information landscape. While the preclinical relevance of the metrics, such as functional connectivity, are yet to be conclusively determined, the additional data provided by an SBI approach can allow future research to create more predictive models to better understand how neural cultures are influenced by pharmacological intervention.

Apart from the specific limitations already discussed, this paper also acknowledges several notable general constraints. Foremost, the *in vitro* neural culture model adopted for this paper is a 2-dimensional monolayer of a single neural cell type. While this allowed rapid generation and iterations in testing different pharmacological compounds at a reasonably high throughput, it is likely that this does not capture the more complex activity which occurs in epilepsy. The generation of 3-dimensional neural organoids in simulated information rich environments has previously been proposed as a method called Organoid Intelligence (OI) [38–40]. An OI approach may offer the ability to determine even more complex interactions between the neural systems and pharmacological interventions. Additionally, future work could also consider generating and evaluating *in vitro* models from donors who display genetically related forms of epilepsy [19, 41, 42]. Another key limitation of this work is that of the limited time window that testing could be undertaken; as previously explored, the *DishBrain* setup allows for restricted testing periods as the heat generated by the MEA system results in evaporation and changes in osmolarity of the cell culture media. These changes eventually result in degradation of cell health and eventual cascades of cell death in the cultures. Future work should aim to use systems that either generate minimal heat or adopt an approach of embedding neural cultures in closed-system perfusion circuits capable of a constant homeostatic control of the cell culturing environment. Further, while the drugs and doses chosen for this study represent established options for the treatment of epilepsy and have been used in similar preclinical drug screens involving NGN2 iPSC-derived neurons [20], a more comprehensive panel of ASMs can be identified, including those with differing mechanisms of action. As such, future work could utilize wider libraries of compounds to gain a deeper insight into the effects of suppression of neuronal excitability on this system. Finally, this work intended to use the previously validated *DishBrain* system to assess the information processing capabilities of a neural system that expressed glutamatergic hyperactivity in the context of epilepsy. While improvements in the cell culture model remain possible for future work, the differences in neural activity between gameplay and rest conditions, and the impact of various pharmacological interventions, were demonstrated across multiple metrics. Of course, it is possible that using enhanced disease models, alternative pharmaceutical interventions, or different stimulation and recording settings might yield different performance outcomes. As such, future work should aim to use more modifiable microphysiological systems capable of trialing different learning environments and rapidly iterating through different stimulation and recording assays.

Ultimately, this study demonstrated that pharmacological modulation of a human iPSC-derived model of glutamatergic hyperexcitability can influence both electrophysiological activity and information processing within a structured, closed-loop environment. By integrating real-time stimulation and feedback through the *DishBrain* system, we captured dynamic responses to ASMs that traditional *in vitro* assays cannot easily resolve. Despite some technical and biological limitations, our results underscore the value of a SBI approach in preclinical research. This method not only enables richer functional assessments of drug effects but also lays the groundwork for more predictive, human-relevant models of neurological disease. As MPS platforms evolve, combining improved cellular models, extended recording times, and adaptive closed-loop paradigms may transform how we evaluate therapeutic interventions—bridging the gap between neuropharmacological modulation, and meaningful improvements in information processing.

## Acknowledgements

The authors wish to gratefully acknowledge the service and support of Dr. Irena (Iska) Carmichael and Monash Micro Imaging (Monash University, 6 Floor Burnet Tower, Alfred Research Alliance, 89 Commercial Rd, Melbourne, Australia, 3004) for their help in obtaining the immunocytochemistry images. Perampanel, as well as generous advice, was provided by Dr. Ben Rollo, Monash University, Melbourne, Australia. Dr. Nicole Kerlero De Rosbo’s assistance in editing the final manuscript was freely offered, gratefully accepted, and invaluable.

## Data Availability Statement

The imaging data in this study can be found in the figures; original high-res versions can be provided upon reasonable request. The raw RNA-Seq data files can be provided upon reasonable request. The electrophysiological datasets generated and analyzed in this study can be found at the following repository: Link to Open Science Framework (OSF) repository.

## Supplementary Figures

**Supp. Fig. 1:**
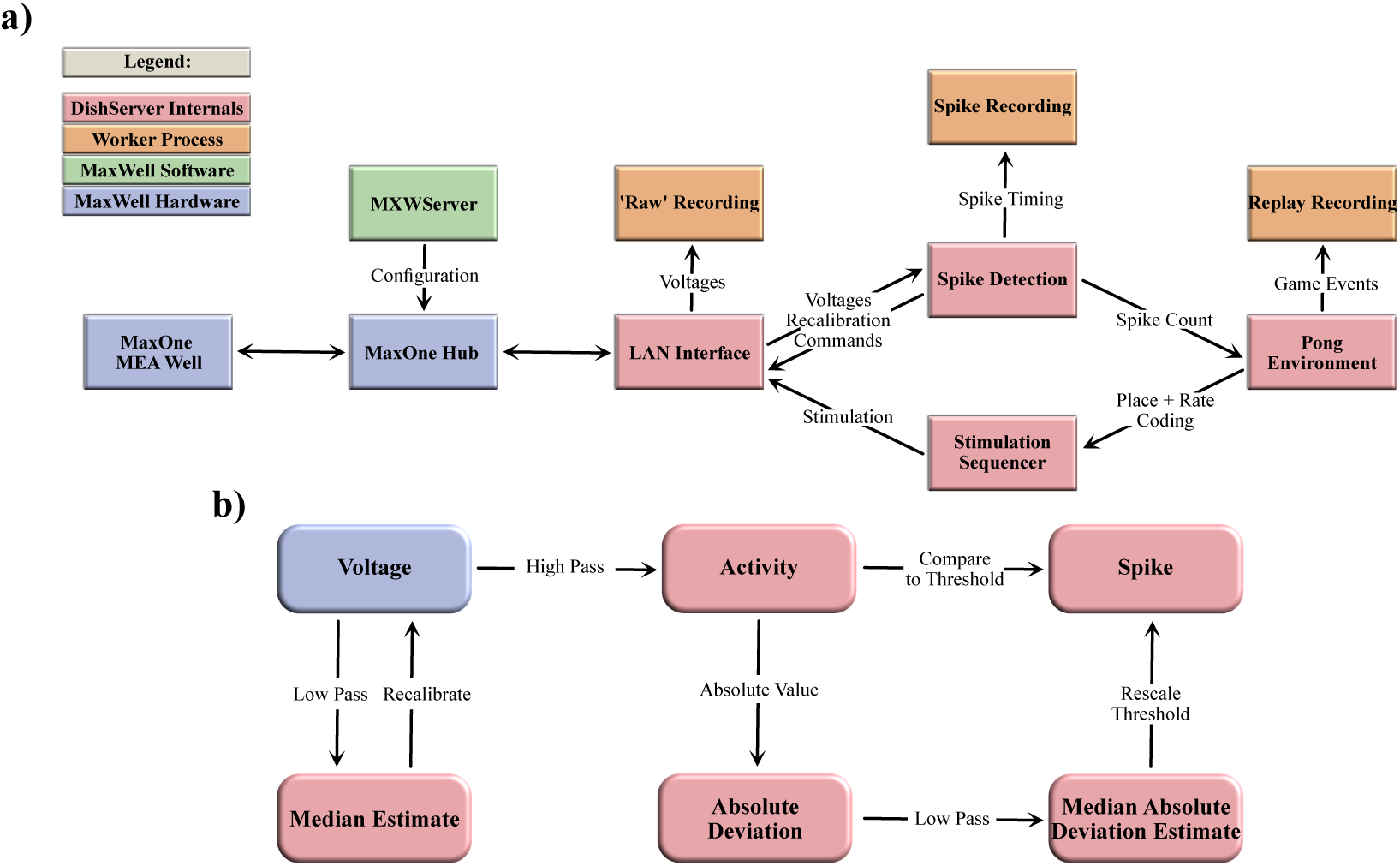
**a, b)** Schematics of software used for DishBrain. **a)** Software components and data flow in the *DishBrain* closed loop system. Voltage samples flow from the MEA to the ‘Pong’ environment, and sensory information flows from the ‘Pong’ environment back to the MEA, forming a closed loop. The blue rectangles mark proprietary pieces of hardware from MaxWell, including the MEA well which may contain a live culture of neurons. The green MXWServer is a piece of software provided by MaxWell which is used to configure the MEA and Hub, using a private API directly over the network. The red rectangles mark components of the ‘DishServer’ program, a high-performance program consisting of four components designed to run asynchronously, despite being run on a single CPU thread. The ‘LAN Interface’ component stores network state, for talking to the Hub, and produces arrays of voltage values for processing. Voltage values are passed to the ‘Spike Detection’ component, which stores feedback values and spike counts, and passes recalibration commands back to the LAN Interface. When the pong environment is ready to run, it updates the state of the paddle based on the spike counts, updates the state of the ball based on its velocity and collision conditions, and reconfigures the stimulation sequencer based on the relative position of the ball and current state of the game. The stimulation sequencer stores and updates indices and countdowns relating to the stimulations it must produce and converts these into commands each time the corresponding countdown reaches zero, which are finally passed back to the LAN Interface, to send to the MEA system, closing the loop. The procedures associated with each component are run one after the other in a simple loop control flow, but the ‘Pong’ environment only moves forward every 200th update, short-circuiting otherwise. Additionally, up to three worker processes are launched in parallel, depending on which parts of the system need to be recorded. They receive data from the main thread via shared memory and write it to file, allowing the main thread to continue processing data without having to hand control to the operating system and back again. **b)** Numeric operations in the real-time spike detection component of the *DishBrain* closed loop system, including multiple IIR filters. Running a virtual environment in a closed loop imposes strict performance requirements, and digital signal processing is the main bottleneck of this system, with close to 42 MB of data to process every second. Simple sequences of IIR digital filters is applied to incoming data, storing multiple arrays of 1024 feedback values in between each sample. First, spikes on the incoming data are detected by applying a high pass filter to determine the deviation of the activity, and comparing that to the MAD, which is itself calculated with a subsequent low pass filter. Then, a low pass filter is applied to the original data to determine whether the MEA hardware needs to be re-calibrated, affecting future samples. This system was able to keep up with the incoming data on a single thread of an Intel Core i7-8809G. Figures adapted from [7].

**Supp. Fig. 2:**
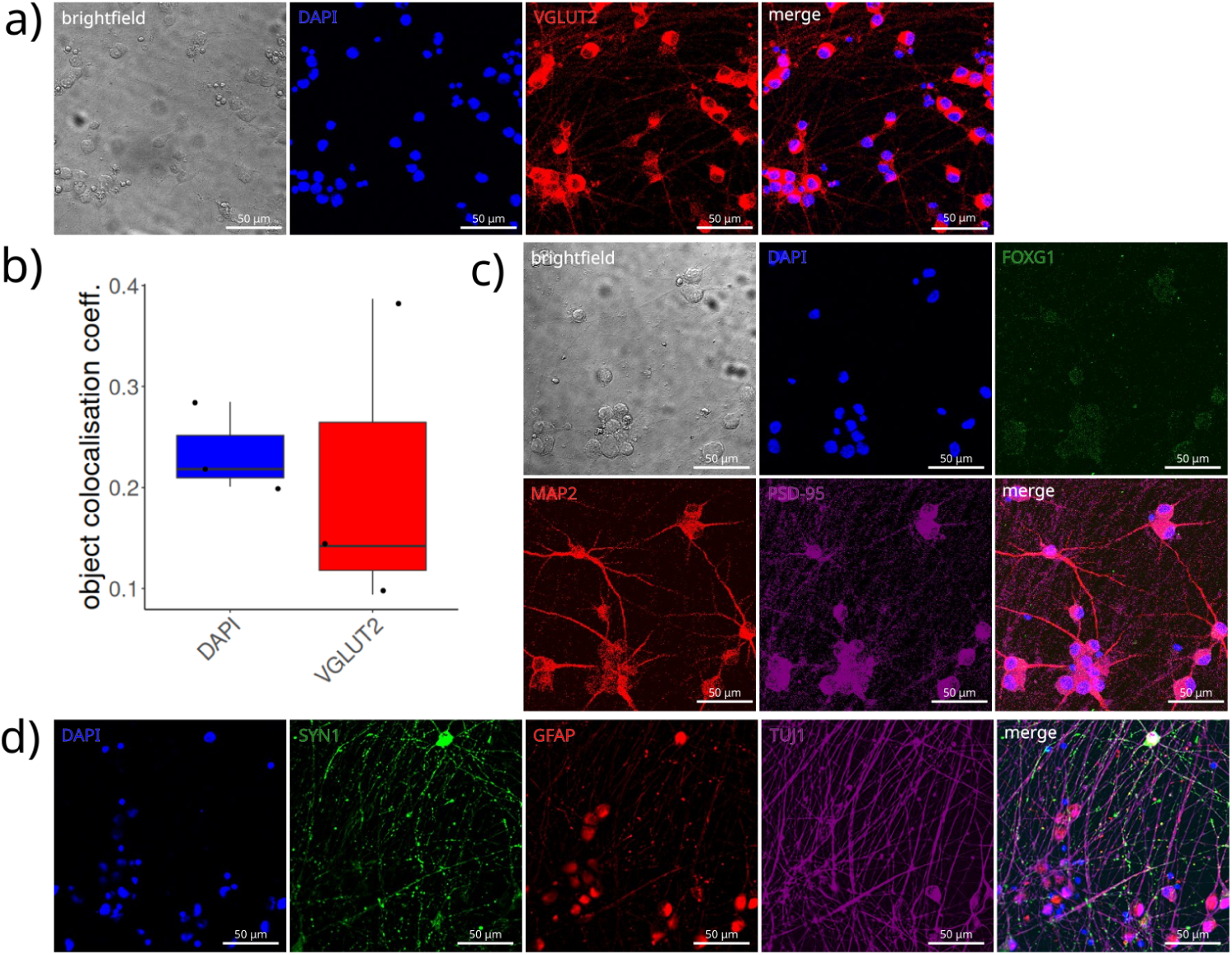
Immunocytochemistry showing separated channels. Same data from Figure 1 b), but showing brightfield field of view and the individual channel fluorescence in addition to the merged images. a) Immunocytochemistry of NGN2 cultures at day 38 showing brightfield, DAPI (blue), and VGLUT2 (SLC17A6; red) labelling. b) Fraction object colocalization overlap of either DAPI or VGLUT2, with objects of the other label. c) Immunocytochemistry in NGN2 cultures showing brightfield, DAPI (blue), FOXG1 (green), MAP2 (red), and PSD-95 (DLG4; magenta) labelling, as well as merged image. d) Immunocytochemistry of NGN2 cultures at day 38 showing DAPI (blue), SYN1 (green), GFAP (red), and TUJ1 (magenta) labelling, as well as merged image. Scale bars in a, c, d); 50 *µ*m.

**Supp. Fig. 3:**
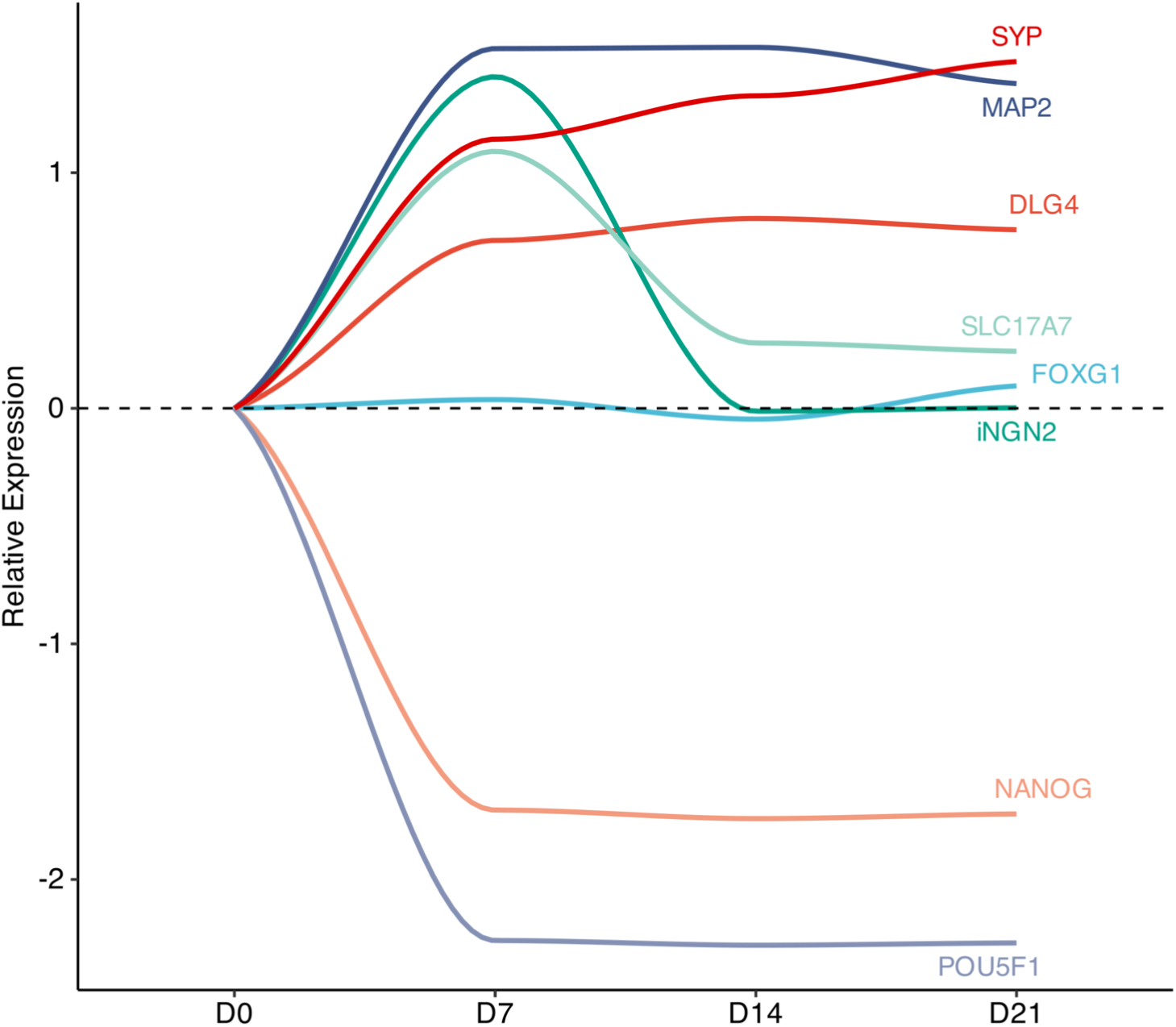
RNA Sequencing results. Fold changes in expression of neural and cortical transcripts before and during induced differentiation. Mean expression is normalized to day 0 (hiPSCs).

**Supp. Fig. 4:**
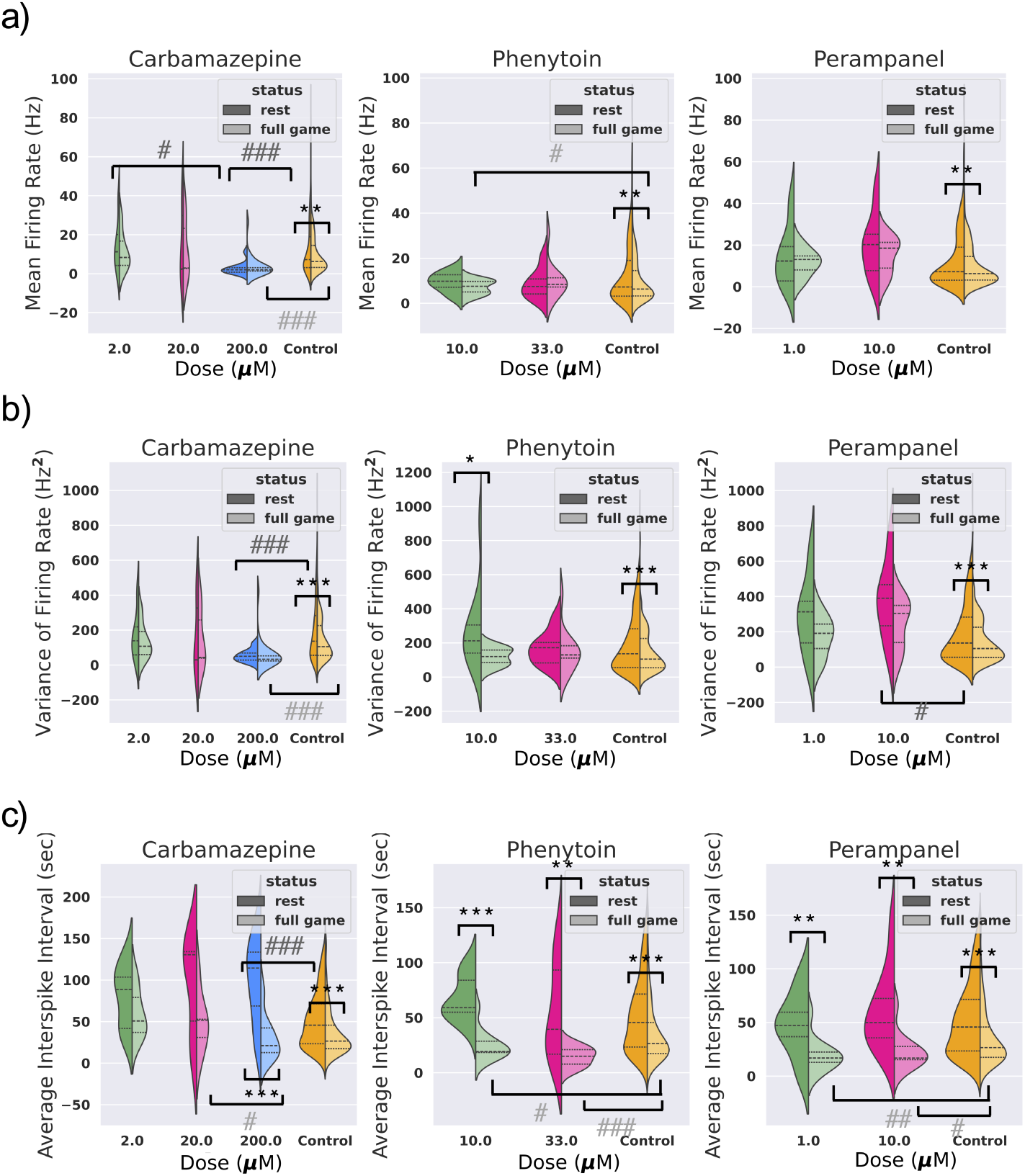
Dose response relationships of ASMs and electrophysiological parameters. a, b, c) Comparison of Mean Firing Rate, Variance of Firing Rate, and Average Interspike Interval (ISI) calculated for different doses of each drug. The sample sizes are n = 12, n = 12; n = 6, n = 6; n = 24, n = 24 (Carbamazepine: rest, full game), n = 12, n = 12, n = 12, n = 12 (Phenytoin: rest, full game), n = 12, n = 11; n = 12, n = 12 (Perampanel: rest, full game) from low dose to high dose respectively. One-way ANOVA test within each group and post-hoc Games Howell test between all the rest and all the full game groups of different doses. # or * indicate *p <* 0.05, ## or ** indicate *p <* 0.01, ### or *** indicate *p <* 0.001.

**Supp. Fig. 5:**
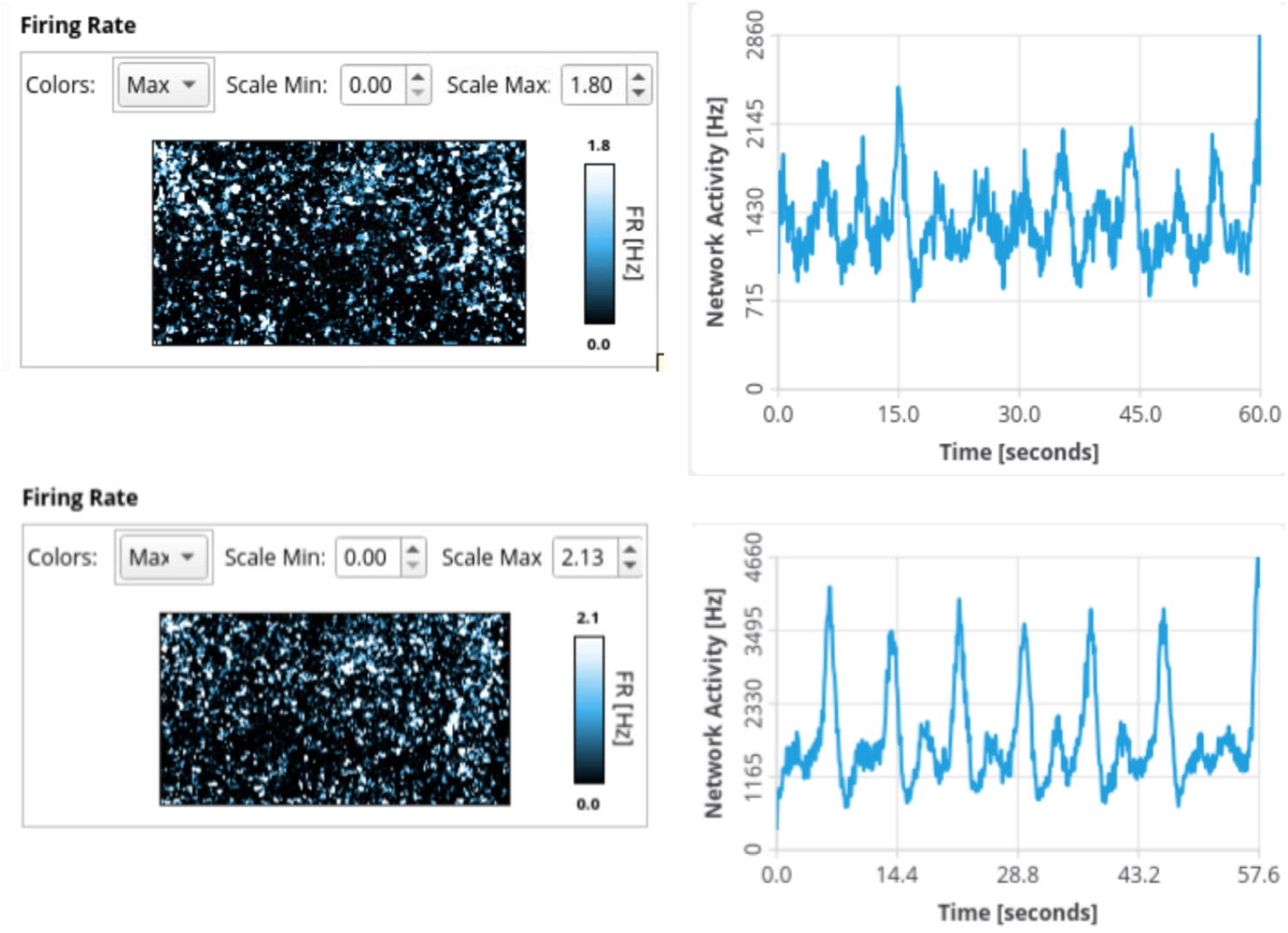
Basic electrophysiological activity of the NGN2 iNeuron network at rest (top panels) and following the addition of the pro-convulsant 4-AP 100 *µ*M (bottom panels). Left panels represent a heat map of where mean firing activity is concentrated on the MEA surface. Right panels show a line plot of total activity in the network over time. Images obtained from MaxLabLive software, version 23.2.2.

**Supp. Fig. 6:**
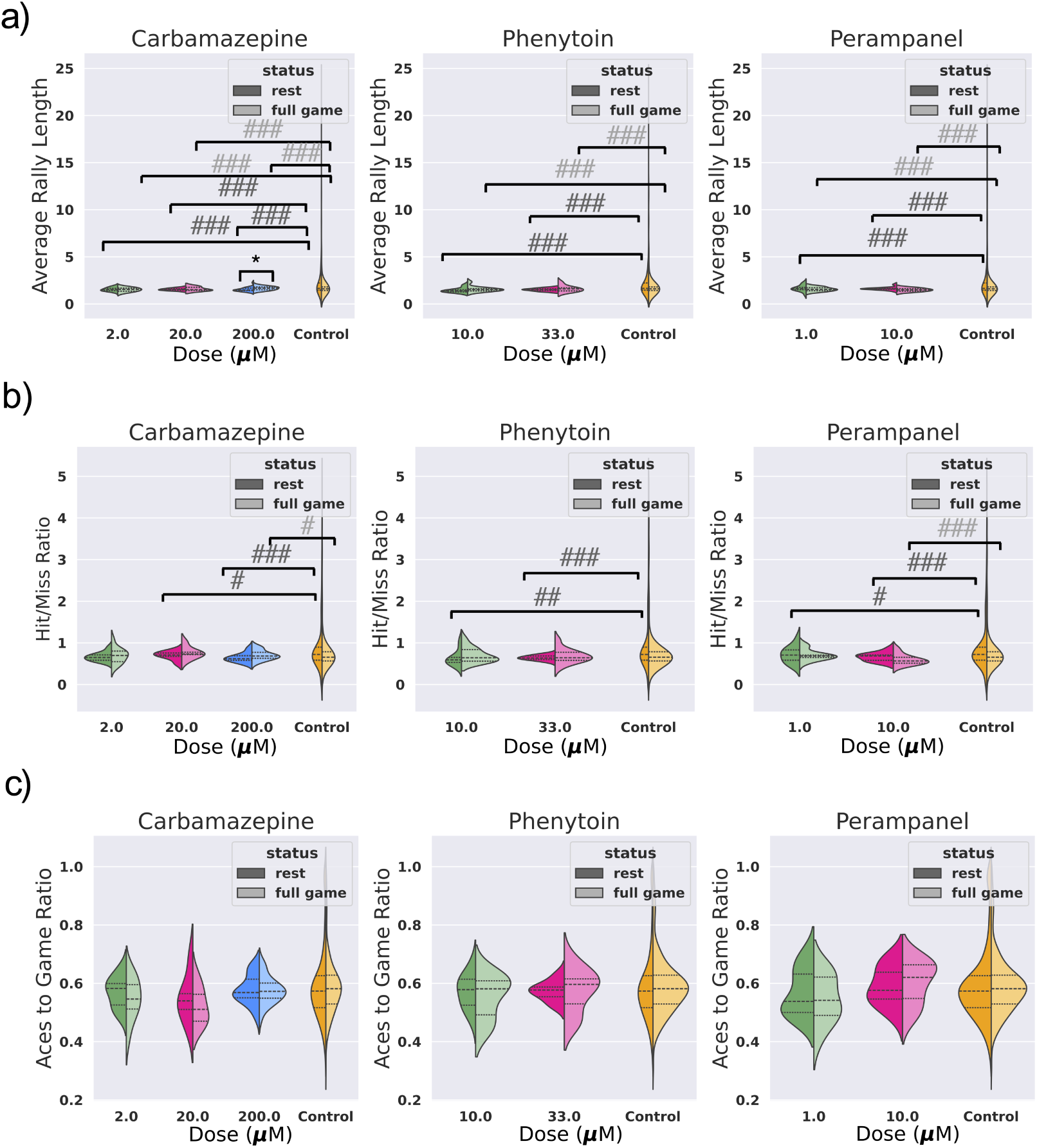
Dose response relationships of ASMs and gameplay parameters. a, b, c) Comparison of Average Rally Length, Hit/Miss Ratio, and Aces to Game Ratio during full game and rest for different doses of each drug. The sample sizes are n = 12, n = 12; n = 6, n = 6; n = 24, n = 24 (Carbamazepine: rest, full game), n = 12, n = 12, n = 12, n = 12 (Phenytoin: rest, full game), n = 12, n = 11; n = 12, n = 12 (Perampanel: rest, full game) from low dose to high dose, respectively. One-way ANOVA test withing each group and post-hoc Games Howell test between all the rest and all the full game groups of different doses. # or * indicate *p <* 0.05, ## or ** indicate *p <* 0.01, ### or *** indicate *p <* 0.001.

**Supp. Fig. 7:**
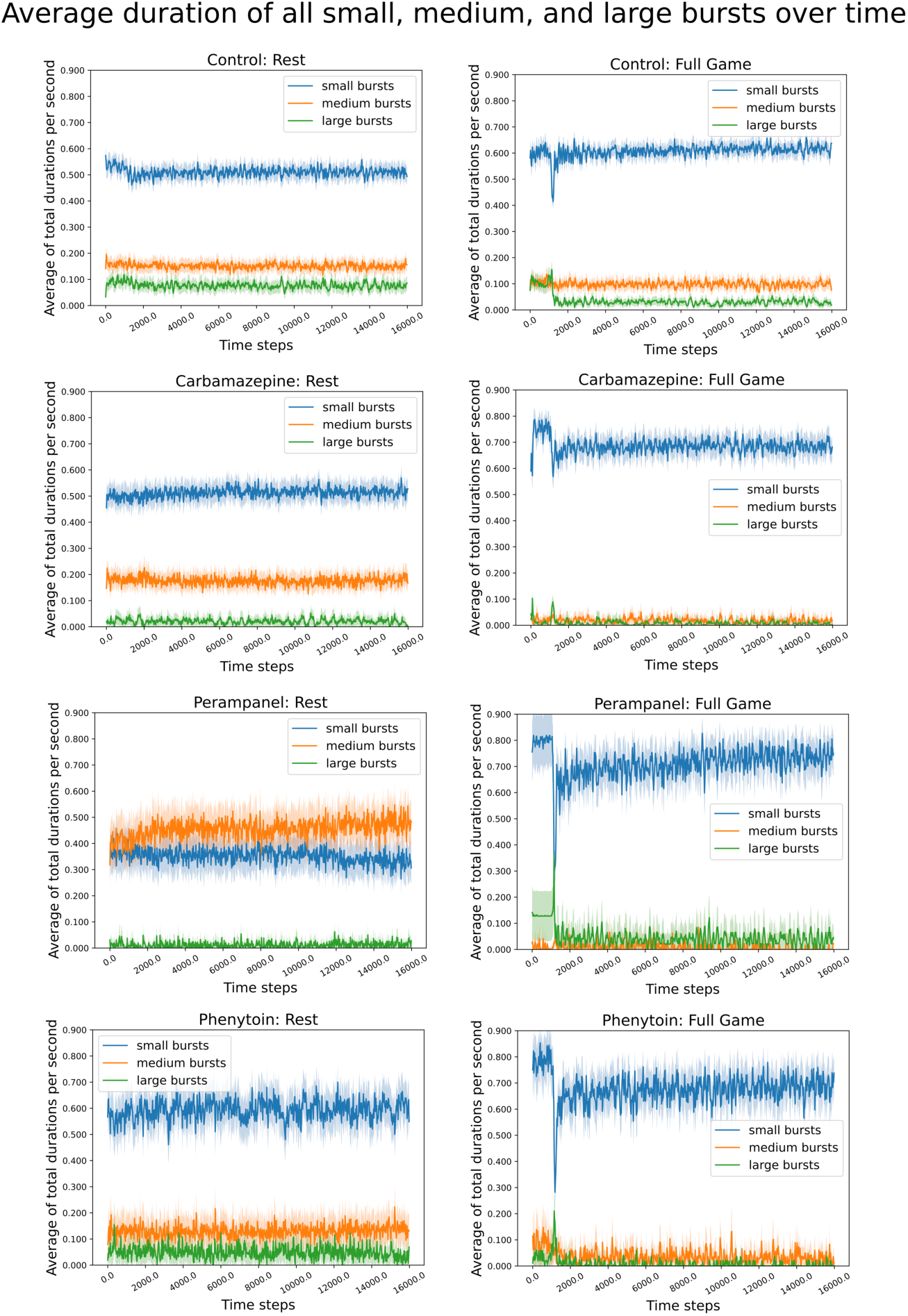
Changes in the proportion of time spent during the occurrence of small, medium, and large bursts per second under control and pharmacological intervention conditions during rest and full game. Lines represent the average values across trials, while shaded areas indicate the 95% confidence intervals.

**Supp. Fig. 8:**
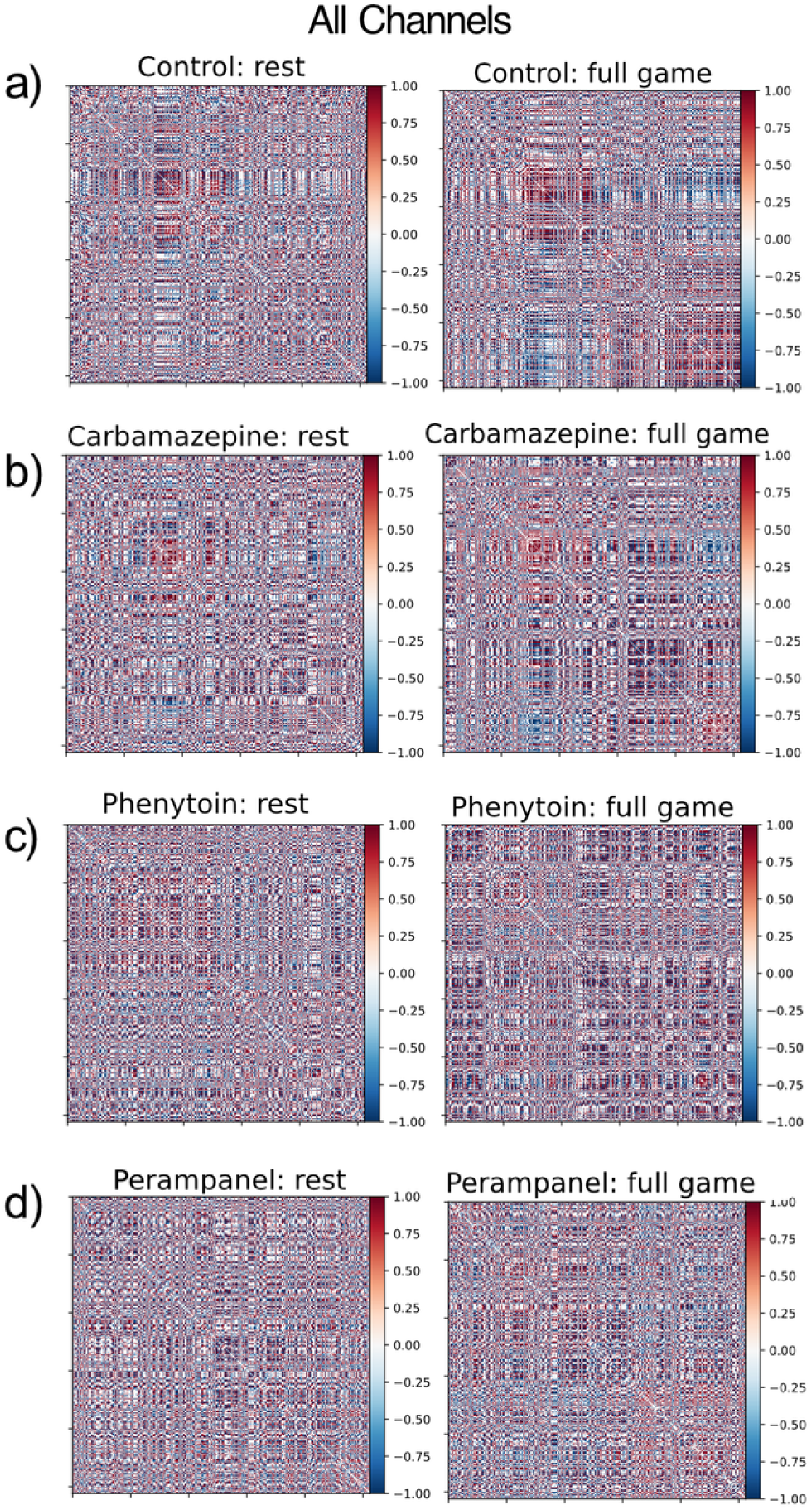
Functional connectivity networks of rest and full game sessions under control and pharmacological interventions. a-d) Heatmaps illustrating the average edge weights between all channels in the Pearson Correlation network constructed from all rest and all full game recordings under control and pharmacological intervention conditions. The sample sizes are n = 42, n = 43 (Carbamazepine: rest, full game), n = 24, n = 24 (Phenytoin: rest, full game), n = 24, n = 24 (Perampanel: rest, full game), n = 354, n = 400 (Control: rest, full game). One-way ANOVA test.

**Supp. Fig. 9:**
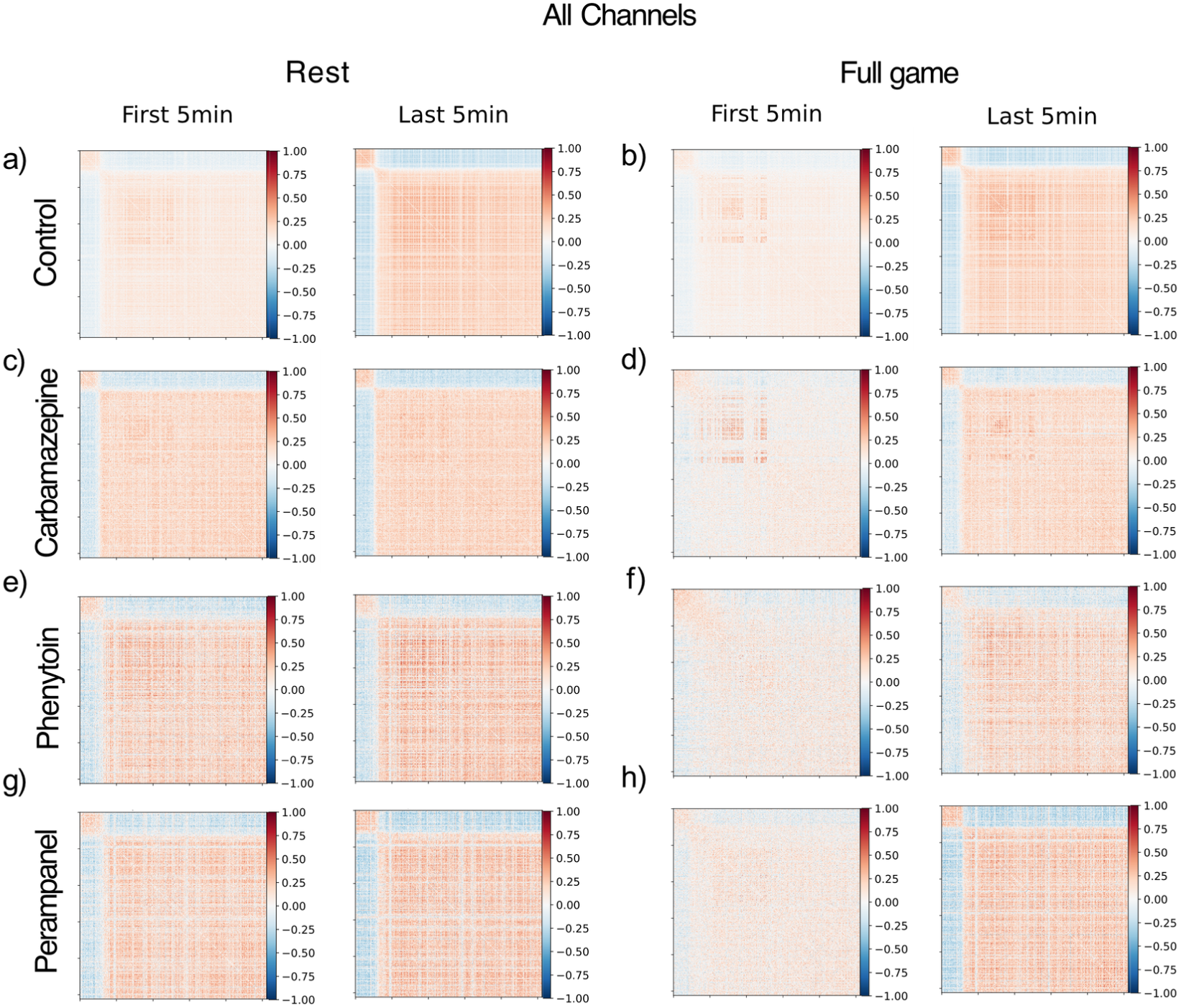
Changes in the functional connectivity networks of rest and full game sessions under control and pharmacological interventions. a-h) Heatmaps illustrate the change in average edge weights between all recorded channels in the Pearson Correlation network constructed from the first and last 5 minutes of all rest and all full game recordings under control and pharmacological intervention conditions.

**Supp. Fig. 10:**
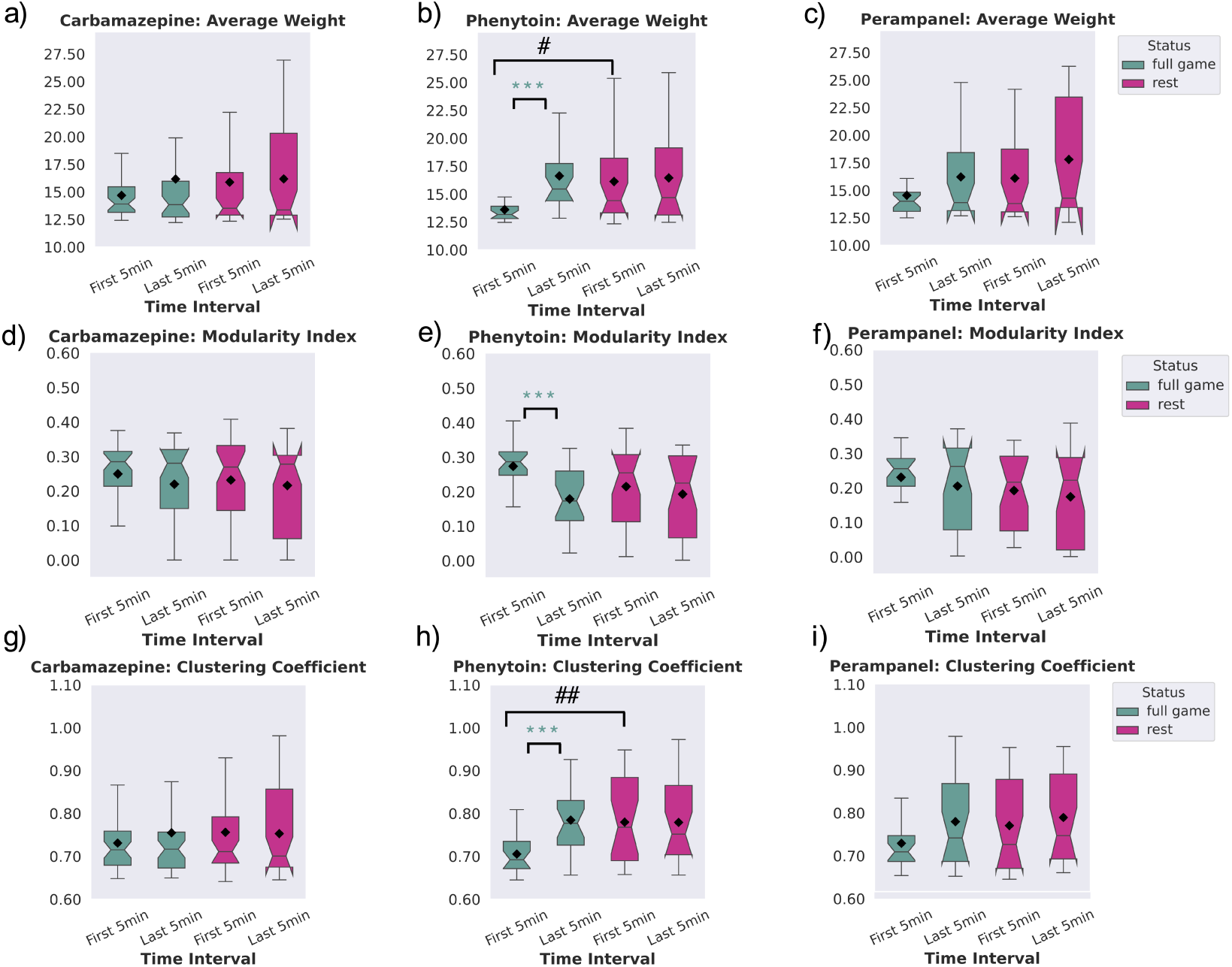
Network summary statistics between the first and last 5 minutes of full game and rest recordings using the 30 representative channels under different drug treatments. a, b, c) Average weight, d, e, f) Modularity Index, and g, h, i) Clustering coefficient values for the first and last 5 minutes of recordings under all pharmacological intervention using the 30 representative channels. One-way ANOVA test withing each group and post-hoc Tukey’s test between all the rest and all the full game groups. # or * indicate *p <* 0.05, ## or ** indicate *p <* 0.01, ### or *** indicate *p <* 0.001.

**Supp. Fig. 11:**
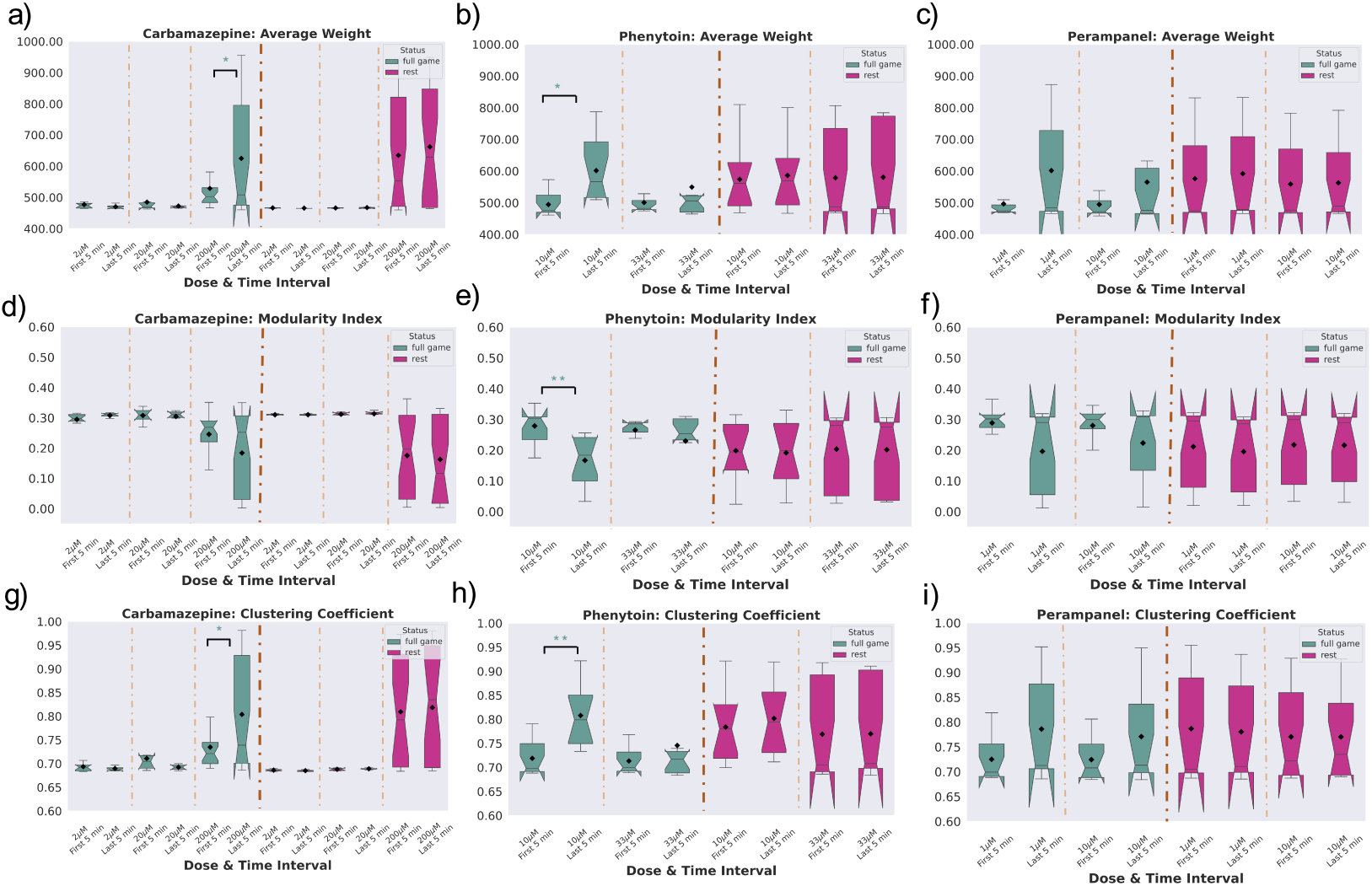
Network summary statistics between the first and last 5 minutes of full game and rest recordings using all of the recorded channels under different doses of drug treatments. a, b, c) Average weight, d, e, f) Modularity Index, and g, h, i) Clustering coefficient values for the first and last 5 minutes of recordings under all doses of the pharmacological interventions using all 1024 recorded channels. One-way ANOVA test withing each group and post-hoc Tukey’s test between all the rest and all the full game groups. * indicates *p <* 0.05, ** indicates *p <* 0.01, *** indicates *p <* 0.001.

## Supplementary Methods

### MEA Setup and Plating

MaxOne Multielectrode Arrays (MEAs; Maxwell Biosystems, AG, Switzerland) was used and is a high-resolution electrophysiology platform featuring 26,000 platinum electrodes arranged over an 8 mm^2^ surface. The MaxOne system is based on complementary meta-oxide-semiconductor (CMOS) technology and allows recording from up to 1024 channels. MEAs were coated with 60 *µ*L Poly-D-Lysine (P6407-5MG, Merck, Australia) dissolved in 1X borate buffer (28341, ThermoFisher, 20x) in ddH_2_0 to a final concentration of 50*µ*g / mL. These chips were incubated for at least 1 hr (or O/N) at 37^◦^C, then washed 3 times with ddH2O and let dry for an hour. Geltrex (A1413302, ThermoFisher, Australia) was then used to coat the PDL-primed MEA. 60 *µ*L of cold Geltrex working solution (made from 250 *µ*L frozen stock aliquot dissolved in 25 mL cold (*<*4^◦^C) DMEM/F-12 media) was added to the MEA surface before returning to the incubator for at least 45 min.

Approximately 2 × 10^5^ neurons and 5 × 10^4^ astrocytes were plated on the MEA after preparation; these numbers were determined after discussions with collaborators and empircal testing. Geltrex solution was aspirated briefly before plating, ensuring that the MEA surface did not dry out. Cells were allowed 1 h to adhere to MEA surface before the well was flooded with cell culture media. The day after plating, cell culture media was fully replaced as detailed in the main methods. Cultures were maintained in a low O_2_ incubator kept at 5% CO_2_, 5% O_2_, 36°C and 80% relative humidity. Every two days, half the media from each well was removed and replaced with fresh media. Media changes always occurred after all recording sessions.

### *DishBrain* platform and electrode configuration

The *DishBrain* platform is configured as a low-latency, real-time MEA control system with on-line spike detection and recording software. The *DishBrain* platform provides on-line spike detection and recording configured as a low-latency, real-time MEA control. The *DishBrain* software runs at 20 kHz and allows recording at an incredibly fine timescale. This setup captured neuronal electrical activity and provided long-term, safe external electrical stimulation through biphasic pulses that elicited action potentials in neurons, as detailed in previous studies [43].

There is the option of recording spikes in binary files, and regardless of recording, they are counted throughout 10 milliseconds (200 samples), at which point the game environment is provided with how many spikes are detected in each electrode in each predefined motor region as described below. Based on which motor region the spikes occurred in, they are interpreted as motor activity, moving the ‘paddle’ up or down in the virtual space. As the ball moves around the play area at a fixed speed and bounces off the edge of the play area and the paddle, the pong game is also updated at every 10 ms interval. Once the ball hits the edge of the play area behind the paddle, one rally of pong has come to an end. The game environment will instead determine which type of feedback to apply at the end of the rally: random, silent, or none. Feedback is also provided when the ball contacts the paddle under the standard stimulus condition. A ‘stimulation sequencer’ module tracks the location of the ball relative to the paddle during each rally and encodes it as stimulation to one of eight stimulation sites. Each time a sample is received from the MEA, the stimulation sequencer is updated 20,000 times a second, and after the previous lot of MEA commands has completed, it constructs a new sequence of MEA commands based on the information it has been configured to transmit based on both place codes and rate codes. The stimulations take the form of a short square bi-phasic pulse that is a positive voltage, then a negative voltage. This pulse sequence is read and applied to the electrode by a Digital to Analog Converter (or DAC) on the MEA.

Alternatively, cells could be recorded at ‘Rest’ in a Gameplay environment where activity was recorded to move the paddle but no stimulation was delivered, with corresponding outcomes still recorded. Using this spontaneous activity alone as a baseline, the Gameplay characteristics of a culture were determined. Low level code for interacting with Maxwell API was written in C to minimize processing latencies-so packet processing latency was typically *<*50 *µ*s. High-level code was written in Python, including configuration setups and general instructions for game settings. A 5 ms spike-to-stim latency was achieved, which was substantially due to MaxOne’s inflexible hardware buffering. Supp. Fig. 1 illustrates a schematic view of Software components and data flow in the *DishBrain* closed loop system.

### Stimulation and Feedback

Specifically, stimulation was applied using a combination of rate coding (4Hz - 40Hz) electrical pulses to communicate the position on the *x*-axis and place coding (on a given electrode that was arranged topographically from an egocentric representation for the culture) to communicate information on the *y*-axis into a predefined bounded two-dimensional sensory area consisting of 8 sensory electrodes to deliver this input information. Three types of input were provided: the sensory stimulation as explained above, or stimulation in response to activity designated as either ‘Predictable’ or ‘Unpredictable’ feedback. Cultures received Unpredictable stimulation when they missed connecting the paddle with the ‘ball’, i.e. when a ‘miss’ occurred. Using a feedback stimulus at a voltage of 150 mV and a frequency of 5 Hz, an unpredictable external stimulus could be added to the system. Random stimulation took place at random sites over the 8 predefined sensory electrodes at random timescales for a period of four seconds, followed by a configurable rest period of four seconds where stimulation paused, then the next rally began. Should no miss occur, the game would continue until either a miss occurred or the timer of 20 minutes expired, which would end the session. In contrast, cultures were exposed to Predictable stimulation when a ‘hit’ was registered - that is, when the ‘paddle’ connected successfully with the ‘ball’. This was delivered across all 8 stimulation electrodes simultaneously at 75mV at 100Hz over 100ms and replaced other sensory information for 100 ms.

The movement of the paddle was controlled by the level of electrophysiological activity measured in a predefined ‘motor area’ of the cultured network, which was collected in real-time. Incoming samples were filtered with a 2nd order high-pass Bessel filter with 100Hz cut-off. The absolute value was smoothed using a 1st order low-pass Bessel filter with a 1 Hz cut-off and the spike threshold is proportional to this smoothed absolute value. A relative activity spike of 6 sigma greater than background noise was then used to define an action potential. Detected action potentials from counterbalanced motor regions were then summed together, where higher activity in a given pair of regions would cause the virtual paddle to move in one direction, while activity in the other regions would result in the inverse movement. Information about ball position relative to the paddle was adjusted in a closed-loop manner with a spike-to-stim latency of approximately 5ms.

The gameplay performance of cell cultures subjected to the simplified pong environment via the *DishBrain* system was assessed. In each episode of the game, the average number of rallies before the ball was missed for the first time was then compared with different deep RL baseline methods. Each recording session of the cultures during gameplay was 15 minutes.

### Description of analysis terms used

**Supp. Table 1:** Firing statistics and game performance metrics with details of their definition.

| Notation | Definition |
| --- | --- |
| Mean Firing Rate (Hz) | The average rate at which recorded channels fire action potentials (spikes) over one second. |
| Variance of Firing Rate (Hz<sup>2</sup>) | Variance of the rate at which recorded channels fire action potentials (spikes) over one second. |
| Average Interspike Interval - ISI (sec) | The average amount of time between consecutive spikes (action potentials) of the channels. |
| Average Rally Length | The average number of times the ball is hit back and forth between the paddle and the wall in a single rally. |
| Hit/Miss Ratio | The ratio of the total accurate hits of the ball to the number of misses. |
| Aces to Game Ratio | The ratio of the number of games missed after the first serve to the total number of games. |

**Supp. Table 2:**
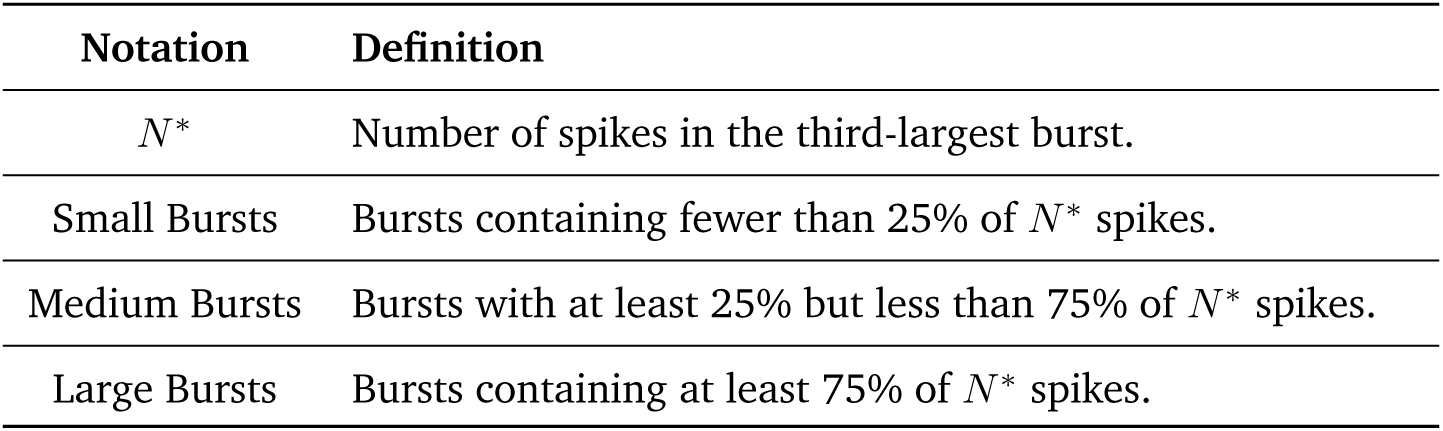
Burst pattern metrics with details of their definition.

| Notation | Definition |
| --- | --- |
| $N^*$ | Number of spikes in the third-largest burst. |
| Small Bursts | Bursts containing fewer than 25% of $N^*$ spikes. |
| Medium Bursts | Bursts with at least 25% but less than 75% of $N^*$ spikes. |
| Large Bursts | Bursts containing at least 75% of $N^*$ spikes. |

**Supp. Table 3:**
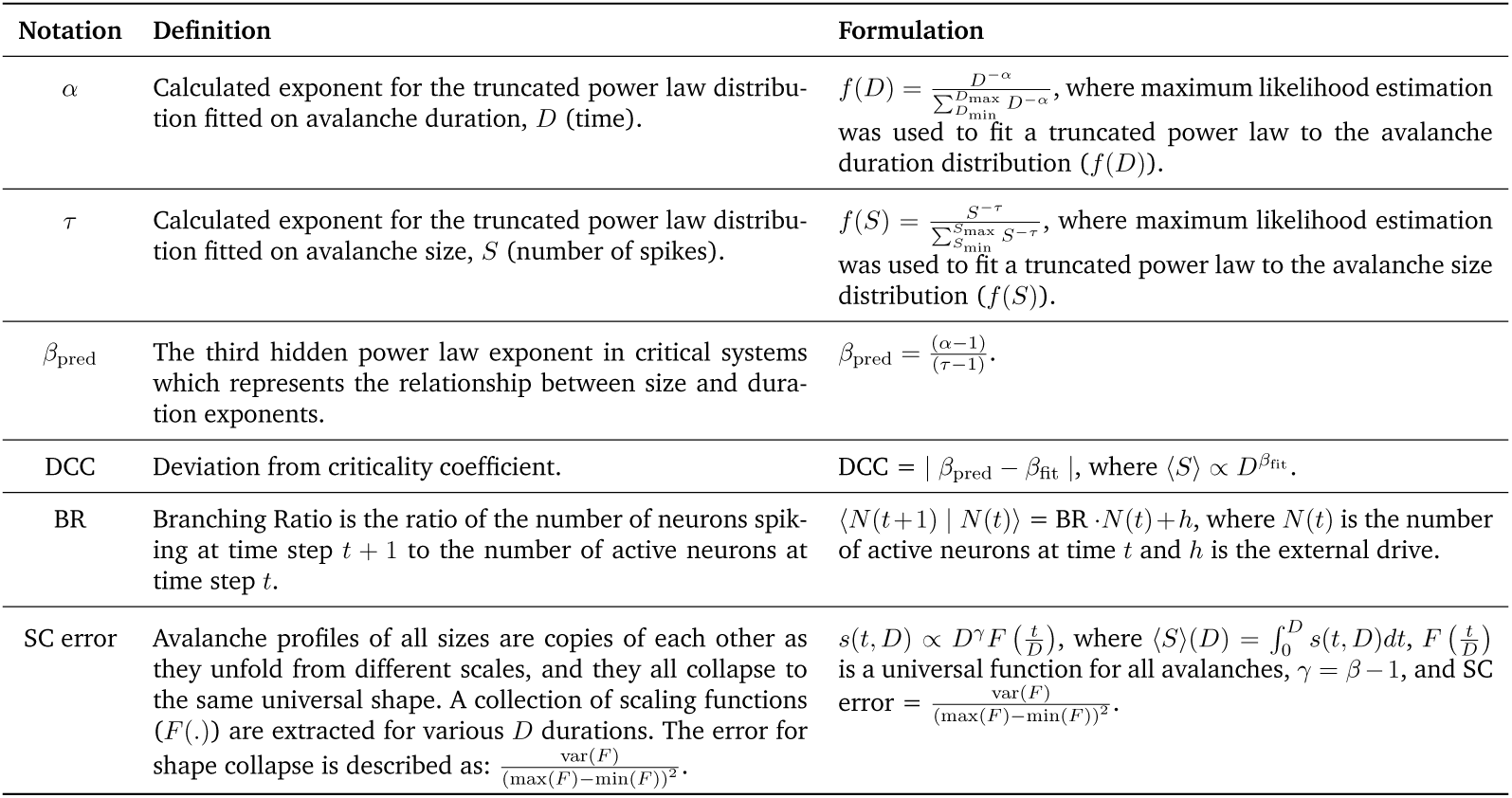
Criticality parameters and metrics with details of their formulation.

### Criticality analysis

Neural systems have been hypothesized to operate near a critical state, characterized by well-defined dynamical properties such as stability of neuronal activity, optimized information storage, and enhanced information transmission [26, 44–47]. The concept of these networks self-organizing near a critical state is supported by decades of research and suggests that the balance between excitatory and inhibitory inputs is crucial for achieving a critical phase transition [37]. Therefore, the criticality analysis helps anticipate such transitions, indicating network robustness and potentially predicting network failure risks, like epileptic seizures [27, 48]. The scale-free dynamics of detected neuronal avalanches, as well as the Deviation from Criticality Coefficient (DCC), Branching Ratio (BR), and Shape Collapse error (SC error) are evaluated to identify whether/when the recordings are tuned near criticality. A lower DCC and SC error and a higher BR (closer to 1) indicate a closer to criticality state. The avalanche analysis is performed in order to study the network in terms of its distance from criticality. The start and stop of an avalanche are determined by crossing a threshold of network activity [46]. An avalanche can be initiated by spikes from any and all neurons within a region of interest. The number of contributing spikes in each avalanche (S) and the total duration of the event (D) are then measured. To demonstrate the distance from criticality in a cultured cortical network, the presence of the following markers were investigated in the dynamics of our data: 1) Power law observables; 2) Exponent relation; 3) Branching ratio parameter; and 4) Scaling function. Certain criteria on these markers are the necessary conditions of a critical regime and meeting those criteria can indicate with a high confidence whether the system lies near a critical point [49, 50].

### Power law observables

A critical system has interacting components (here, neurons) that show some fluctuation in their activity while also maintaining a level of correlation between their individual activities (here, individual spiking). Criticality implies that the system is defined by scale free dynamics and that events in both the spatial and temporal domains obey power laws [51–53]. For the networks considered here, events are contiguous cascades of spiking activity, rather than limited local bursts of spiking activity or huge network-wide spiking events. These contiguous cascades of spiking activity are called neuronal avalanches. To investigate this property in our system, binary spike trains of each neuron’s activity were utilized. The whole duration of each recording session was discretized to 50 ms bins. The sum of all cells’ activities in each time bin was used as the network activity. Next, a threshold of 40% of the median spiking activity in the network among all time bins was introduced. The start and end points of an avalanche were defined as the time points when the network activity crossed this threshold value from below and then above [29]. Our results were statistically robust across a range of activity thresholds between 30% and 70%. The size of an avalanche, S, is the total number of spikes during the avalanche. The avalanche duration, D, is the time between threshold crossings. Similar to Ma et al. [46], maximum likelihood estimation was used to fit a truncated power law to the avalanche size distribution:

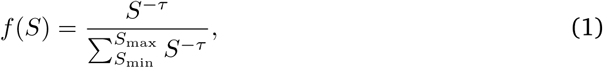

where *τ* is the power law exponent corresponding to avalanche sizes. For a neuronal recording session in which *N_A_* avalanches are detected, the fitting process to obtain the above equation is the following iterative procedure [54]:

1. Find the maximum observed avalanche size *S*_max_.
2. Evaluate the three different power law exponents, *τ*, for the 3 smallest avalanche sizes observed, *S*_min_.
3. Calculate the *Kolmogorov-Smirnov* (KS) test for this estimation to determine the goodness-of-fit between the fitted power law and the empirical distribution.
4. Among the obtained KS values, choose the smallest one, together with the corresponding *τ* and *S*_min_ values.
5. Complete the estimation if 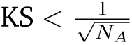 or otherwise repeat steps 2 to 5 with *S*_max_ reduced by 1 until this condition is met.

Steps 3 to 5 are necessary to ensure the data distribution indeed comes from a power law rather than another candidate heavy tailed distribution, such as log normal and stretched exponential forms [55]. Applying the exact same procedure to the set of *D* avalanche events, the corresponding power law exponent of *α* was calculated for the entire avalanche duration distribution.

### Exponent relation and Deviation from Criticality Coefficient (DCC)

In critical systems, there is another exponent relationship between the power law parameters (*α* and *τ*) and the exponent of mean avalanche sizes (⟨*S*⟩), given their duration, *D* [56]. We first find this third power law exponent of the system, *β*, from the experimental data using linear regression given the following exponent relation is present in a critical system:

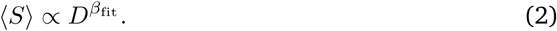

This third power law exponent also relates the size and duration distributions of the avalanches and is predicted by:

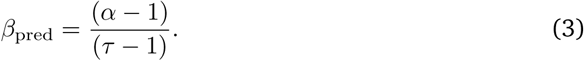

Comparing the fitted value from the empirical data (*β*_fit_) and its estimation using *α* and *τ* exponents (*β*_pred_), a new measure is derived to evaluate the *Deviation from Criticality Coefficient* (DCC), parameterised as *d_CC_*:

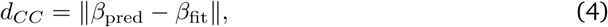

where *β*_pred_ and *β*_fit_ are the predicted and fitted values of *β* respectively. Consequently, a smaller DCC value indicates a more accurately fit power law distribution to the empirical data.

### Branching ratio

The branching ratio is defined as the ratio of the number of units (neurons) active (spiking) at time step *t* + 1 to the number of active units (neurons) at time step *t*. Since a critical regime is naturally balanced and avoids runaway gains, the critical branching ratio is 1. Consequently, on average, network activity neither saturates nor dampens over time. Suppose that *N* active neurons are detected in total and the number of active neurons in each time step *t* is defined by *N* (*t*). A fixed branching ratio of *m*, gives:

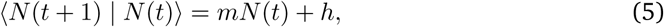

where ⟨|⟩ is the conditional expectation and *h* is the mean rate of external drive. The activity decreases if *m <* 1, whereas it grows exponentially if *m >* 1, meaning that *m* = 1 separates these two regimens and represents a critical dynamic point. A precise prediction of *m* helps to assess the risk that *N* (*t*) will develop large and devastating avalanches of events such as epileptic seizures.

Under the circumstances when the full activity *N* (*t*) is known, *m* can be conventionally estimated using linear regression. Nevertheless, when using subsampling, when only a fraction of neurons in a neuronal network are sampled, this conventional method will be biased to some extent. The bias vanishes only if all units are sampled, because it is inherent to sub-sampling and cannot be overcome by obtaining longer recordings. Instead, inspired by the method introduced in [57] the subsampled activity *n*(*t*) is utilized, where the fraction of recorded units to all cells is defined as a constant *µ*. *n*(*t*) here is a random variable whose expectation is proportional to the real *N* (*t*) and ⟨*n*(*t*) | *N* (*t*)⟩ = *µN* (*t*)+*ξ*, where *µ* and *ξ* are constants. The bias value for the conventional linear estimator can now be calculated as:

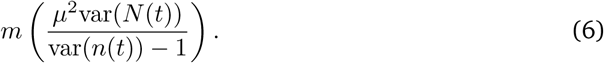

To overcome this subsampling bias, the method introduced by Wilting and Priesemann [57] was utilized. Instead of directly using the biased regression of activity at time *t* and *t* + 1, multiple linear regressions of activity between times *t* and *t* + *k* were performed with different time lags *k* = 1*, …, k*_max_. Each of these *k* values returns a regression coefficient *r_k_* with *r*_1_ being equal to the result of a conventional estimator of *m*. With subsampling, all these regression slopes are biased by the same factor 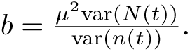 In these circumstances, instead of the exponential relation *r_k_* = *m^k^* which is expected under full sampling, the equations generalizes to:

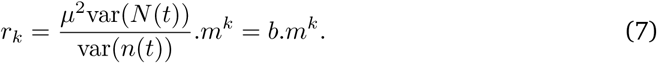

Having multiple calculated *r_k_* values, both *b* and *m* are estimated, which are constant for all *k*.

### Scaling function

Another feature of critical dynamics is that avalanche shapes show fractal properties and all avalanche profiles of different sizes are scaled versions of the universal same shape. According to [56], the value of *β* obtained from the exponent relation analysis can be used to calculate a scaling function for the avalanche shapes. For any given avalanche duration *D*, the average number of neurons firing at time *t* (within *D* seconds) is defined by *s*(*t, D*). The following relations hold in this system:

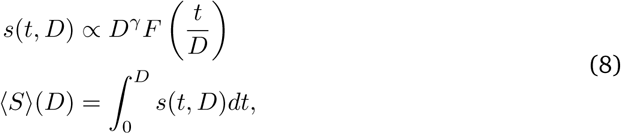

where 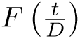 is a universal function for all avalanches and *γ* = *β* − 1. Hence in this process, an initial *β* is used to predict *γ* and using this *γ* and the first term in Equation 8, 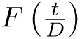 is obtained as 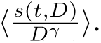 Here ⟨.⟩ denotes the average over all avalanches with duration *D*. A collection of 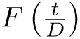 functions are extracted for various *D* durations. The error for shape collapse is described as:

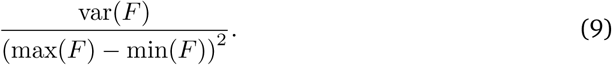

Repeating this process with various values for *β*, the exponent that produces the smallest shape collapse error in Equation 9 is selected as the final scaling factor. In principle, we expect to obtain similar (if not the same) *β* values from this analysis and the estimates in Section 1. The NCC toolbox in MATLAB [58] was utilized to perform shape collapse on data. This shape collapse error is expected to be minimized under critical conditions. For shape collapse, avalanches with durations from 4 to 20 bins (200 to 1,000 ms) were considered. Across the time course of our recordings, there were not enough avalanches to conduct meaningful shape collapse analysis, beyond these cutoffs.

### Burst pattern analysis

Inspired by the methods and metrics for the classification of bursts (or avalanches) introduced in Wagenaar et al. [25], we extracted the following quantitative details from the recordings of each *in vitro* culture to study the size distributions of bursts. Below is a brief explanation of each calculated criteria to identify the small, medium, and large bursts: The range of burst sizes varied among recordings. In every recording from a specific culture, let *N* ∗ denote the number of spikes in the third-largest burst. Bursts containing at least 75% of *N* ∗ spikes were classified as large, while those with at least 25% but less than 75% of *N* ∗ spikes were classified as medium. Bursts containing fewer than 25% of *N* ∗ spikes were labeled as small.

